# Spatiotemporal Dynamics of fMRI Signal Changes Induced by High Concentration Normobaric Oxygen Inhalation

**DOI:** 10.1101/2025.07.29.667573

**Authors:** Yu-Shiang Su, Chien-Te Wu, Joshua Oon Soo Goh, Jerome N. Sanes, Zenas C. Chao

## Abstract

While it is well known that oxygen supports the brain’s metabolic demands, it remains unclear how increased oxygen concentration influences intrinsic neural activity over time and across brain regions. Using resting-state functional magnetic resonance imaging (fMRI), we examined the dynamic responses to high-concentration normobaric oxygen across distinct phases of exposure and withdrawal. We revealed three patterns in the BOLD signals: increased activation during inhalation, an undershoot following immediate oxygen withdrawal, and reactivation even without continued oxygen. These responses were most pronounced in the default mode network (DMN), but also exhibited spatiotemporally heterogeneous patterns across the brain, a map we term Brain Oxygen Sensitivity Topography (BOST). Functional connectivity analyses further revealed increased between-network connectivity during inhalation and enhanced within-network connectivity in the DMN during the aftereffect. This spatiotemporal heterogeneity and transient network reorganization suggests that distinct physiological processes are engaged at each phase, enabling us to predict how different oxygen protocols will enhance specific cognitive functions.

## Introduction

Oxygen is needed for the metabolic requirements of cells in the brain, a process precisely regulated in response to changing energy demands and oxygen availability (1). Perturbations in oxygen intake may alter energy metabolism pathways in neurons (2, 3), affecting the production of ATP that is critical for restoring membrane potentials and supporting synaptic activity. These metabolic changes can influence neuronal excitability and information processing, ultimately shaping behavior. However, how the brain dynamically responds to changes in oxygen availability remains poorly understood. Specifically, it remains unclear which brain regions are involved, how rapidly these changes unfold, and whether they follow consistent spatial or temporal patterns. Moreover, how these neural responses translate into changes in cognition or behavior is not well characterized. Clarifying these mechanisms is essential for understanding how oxygen influences brain function and for guiding future efforts to examine cognitive outcomes through oxygen modulation.

Evidence linking altered oxygen levels to changes in cognition and behavior are supported by a meta-analysis showing that normobaric high-concentration oxygen inhalation with inspired oxygen fractions above 30%, which results in inspired oxygen pressures between 210 and 710 mmHg, has modest but predominantly positive effects on various aspects of cognitive performance (4), including improved accuracy in episodic memory recall and working memory, as well as faster response times in detecting infrequent stimuli. We note that although higher oxygen levels achieved in hyperbaric environments (typically more than one and less than three ATA) has shown benefits for neurological conditions such as traumatic brain injury, stroke and vascular dementias, its application also entails risks, including oxygen toxicity, barotrauma, and pulmonary complications (5–7). Thus, we focused on inhalation of normobaric high-concentration oxygen levels, which has minimal risks, and its effects on brain dynamics and associated cognitive functions. Notably, it is important to recognize that cognitive enhancements have been reported both during exposure to normobaric high-concentration oxygen (e.g. Chung et al., 2007 (8)) and after its withdrawal (e.g. Scholey et al., 1999 (9)), suggesting that the timing of exposure plays a critical role. Moreover, the timing of oxygen exposure appears to differentially influence performance across cognitive domains (4). For instance, memory performance, which is implicated in medial temporal lobe function, shows no significant difference whether assessed during inhalation or after withdrawal. In contrast, visual matching tasks, which are supported by occipital and parietal regions involved in visual processing and attention, show greater enhancement during inhalation than after withdrawal. These findings suggest that the modulation of neuronal excitability by oxygen is both time-dependent and region-specific. Despite these observations, no study to date has systematically examined how different brain regions dynamically respond to different temporal phases of highconcentration oxygen administration, such as during inhalation and following withdrawal. A systematic investigation of these time-dependent and region-specific changes is needed to identify when and where oxygen has its strongest effects on the brain, and how these changes support cognitive functions.

To comprehensively map neural responses to high-concentration oxygen, we used functional magnetic resonance imaging (fMRI) to develop a “Brain Oxygen Sensitivity Topography” (BOST). This approach systematically examines the temporal dynamics both during exposure to high-concentration oxygen and following its withdrawal. Several electrophysiological studies have shown that oxygen inhalation can reduce spectral power in alpha, beta, and gamma bands while increasing activity in the theta and delta bands (10–12). However, electrophysiological recordings have limited spatial resolution, approximately 1 cm^2^, making it difficult to determine exactly how oxygen intake modulates activity across specific brain networks. This limitation can be somewhat overcome by using fMRI, whose higher spatial resolution allows for a more detailed investigation of these effects.

A key challenge in using fMRI for BOST is that blood oxygen level dependent (BOLD) signals primarily reflect the oxygenation state of hemoglobin (13, 14). Therefore, it is crucial to distinguish changes in oxygenation due to local neural activation from those related to globally induced by extra oxygen availability. To solve this potential conflict, we considered the mechanistic process for each contribution. Neural activation consumes more oxygen from local capillary networks that induced local increases in blood flow and raised venous oxyhemoglobin levels (15, 16), leading to local BOLD signal enhancements. By contrast, hyperoxia globally increases dissolved plasma oxygen that reduces the dissociation rate of bound oxygen from the hemoglobin throughout the brain, preserving more oxyhemoglobin in the veins (17–19), which then yields widespread BOLD signal increases (20). Thus, by controlling for fluctuations in global venous oxygenation, the confounding effects of vascular signals on BOLD fluctuations can be identified and removed, allowing localized BOLD changes to more accurately reflect focal neural activity. Based on these facts, we implemented a signal decomposition method to regress out non-neurally driven components including vascular artifacts. After removing the influence of global vascular oxygenation on BOLD signals, we evaluated the temporal dynamic effects and regional responses to high-concentration oxygen inhalation.

We found that the default mode network (DMN) most prominently exhibited enhanced activation during inhalation of highconcentration oxygen. Notably, this enhanced activation dissipated immediately upon oxygen withdrawal but increases again after approximately 5 min. Occipital visual network consistently showed weaker responses, and other brain regions displayed more varied temporal patterns. Functional connectivity analysis revealed increased connectivity between brain networks during oxygen inhalation. By contrast, 5 min after oxygen withdrawal, both connectivity between networks and connectivity within the DMN strengthened, suggesting that oxygen levels can modulate neural network dynamic, that in turn influences both information integration and modular specificity at different phases. These findings highlight the spatiotemporal heterogeneity of brain responses to oxygen inhalation and withdrawal. By leveraging this variability in a functional decoding analysis, we might be able to predict which cognitive functions are likely to be influenced by different oxygen protocols.

## Results

We recruited 22 participants (mean age of 23.34 yr, SD = 2.49, seven females) without neurological, psychiatric, respiratory or cardiac disorders, according to their self-report. Written informed consent was obtained from all participants, and the study was approved by the Research Ethics Committee of the University of Tokyo. Each participant completed two 20-min restingstate fMRI runs, each involving alternating inhalation of normal air and high-concentration normobaric oxygen (Fig. 1A). Air and oxygen were administered through a nasal cannula at a flow rate of 5 l/min (see more details in Methods), resulting in participants inhaling high-concentration oxygen with an inspiratory oxygen fraction (FiO2) of approximately 40% (21), nearly twice the oxygen level of ambient air. This concentration is a common setting for normobaric oxygen delivery via nasal cannula, as it avoids discomfort to the nasal mucosa (21), and has been associated with improvements in cognitive performance (8). During each run (Fig. 1B), participants were instructed to fixate on a central dot displayed on a monitor and breathe through their nose for the entire 20-min scan. Each scan run was divided into four 5-min phases: (1) P1: 0–5 min, the baseline while inhaling normal air; (2) P2: 5–10 min, inhaling oxygen; (3) P3: 10–15 min: switching back to normal air; (4) P4: 15–20 min: continuing with normal air to examine potential post-oxygen aftereffects.

**Fig. 1.**
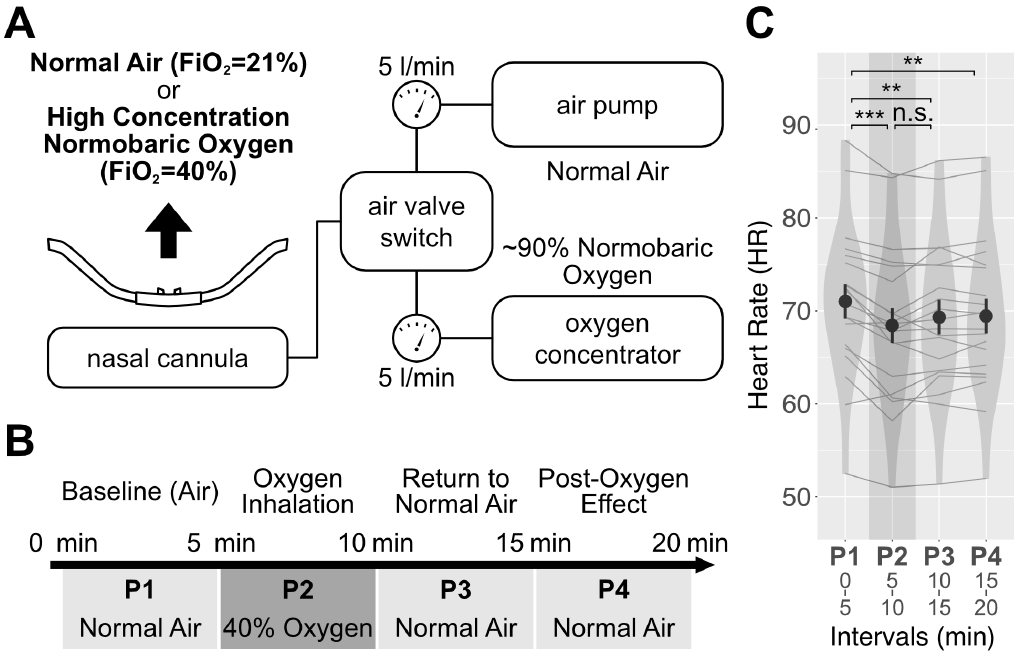
Oxygen delivery setup, experimental protocol, and heart rate changes. (A) A valve switch system was used to control the delivery of air and high-concentration oxygen (40% inspiratory oxygen fraction) at a constant flow rate of 5 l/min through a nasal cannula. This setup enabled phase-specific transitions between normal air (21% oxygen) and high-concentration oxygen in accordance with the experimental protocol. (B) Participants underwent two resting-state fMRI runs. Each run lasted 20 min and included a 5-min period of high-concentration oxygen inhalation. The protocol began with 5-min of normal air, followed by 5-min of high-concentration oxygen, and concluded with 10 min of normal air. To analyze the effects over time, the run was divided into four 5-min phases: P1 (baseline, normal air), P2 (oxygen inhalation), P3 (immediate oxygen withdrawal, return to normal air), and P4 (post-oxygen normal air). (C) Heart rate changes across the four 5-min phases. Oxygen was delivered during P2 (shaded area). Dots and error bars represent the mean and standard error of heart rate. Each line shows an individual participant’s average heart rate trajectory across the two runs over time. We conducted a repeated measures ANOVA with phase and run as within-subject factors, followed by post-hoc comparisons, as illustrated in the figure (***: p < 0.001; **: p < 0.01; n.s.: not significant). All p-values were adjusted for multiple comparisons using the Tukey method.

Cardiac and respiratory signals were recorded throughout the resting-state scans. Due to hardware issues, cardiac data from two participants and respiratory data from four participants were invalid. Several standard physiological metrics were derived (see details in Methods) from the valid recordings. We used a repeated-measures ANOVA to analyze metric changes across phases (P1-P4), resting-state runs (i.e. the first or second 20-min run) were included as a secondary factor to control for potential time-related drift over the course of the experiment. Among these, only heart rate (HR) showed a significant main effect of phases (*F*_3,57_ = 10.65, p < 0.001, *η*^2^ = 0.36) without a significant effect of resting-state runs (*F*_1,19_ = 1.39, p = 0.25, *η*^2^ = 0.07), indicating notable changes between phases (Fig. 1C; see Supplementary Result 1 and Supplementary Fig. 1 for other metrics). Post-hoc pairwise comparisons with a Tukey adjustment revealed a decrease in HR during high-concentration oxygen inhalation (5–10 min) compared to the baseline (0–5 min) (P2 > P1: *t*_57_ = −5.55, p < 0.001, *η*^2^ = 0.35). HR showed a trend toward increasing after returning to normal air; however, this trend was not significant (10–15 min) (P3 > P2: *t*_57_ = 1.91, p = 0.234, *η*^2^ = 0.06). Notably, HR did not return to baseline levels after oxygen withdrawal, remaining lower during 10–15 min (P3 > P1: *t*_57_ = −3.64, p = 0.003, *η*^2^ = 0.19) and 15–20 min (P4 > P1: *t*_57_ = 3.38, p = 0.007, *η*^2^ = 0.17). These results confirmed that high concentration oxygen inhalation significantly affected HR, consistent with previous literature (8, 10, 22–24). Moreover, we found the persistence of reduced HR even after oxygen withdrawal, which suggests a lasting physiological effect within the 10-minute timeframe. Previous studies have reported that HR typically returns to baseline within 10 minutes after oxygen withdrawal (25). Although experimental conditions may differ, it is likely that HR in our study would have recovered beyond this window.

### Vascular Contributions on BOLD Signals During High-concentration Oxygen Inhalation

We initiated our analysis on fMRI data by examining changes in BOLD signals while participants were inhaling high-concentration oxygen. Functional MR images from all participants underwent minimal preprocessing, and voxel-wise signals were scaled to reflect the percentage of signal changes (see details in Methods). The average BOLD signals during high-concentration oxygen inhalation phase (P2) were extracted and compared to those during baseline phase (P1). The resulting voxel-wise map (Fig. 2A) revealed prominent signal changes along the contours of the gray matter surface and the third ventricle. These regions align with major cerebral veins and arteries in the brain. In the sagittal plane, the greatest signal changes, as expected, occurred in the superior sagittal sinus, straight sinus, and great cerebral vein. Additionally, we observed enhanced signal changes along the insula and midbrain, aligning with the vascularization of the middle and posterior cerebral arteries.

**Fig. 2.**
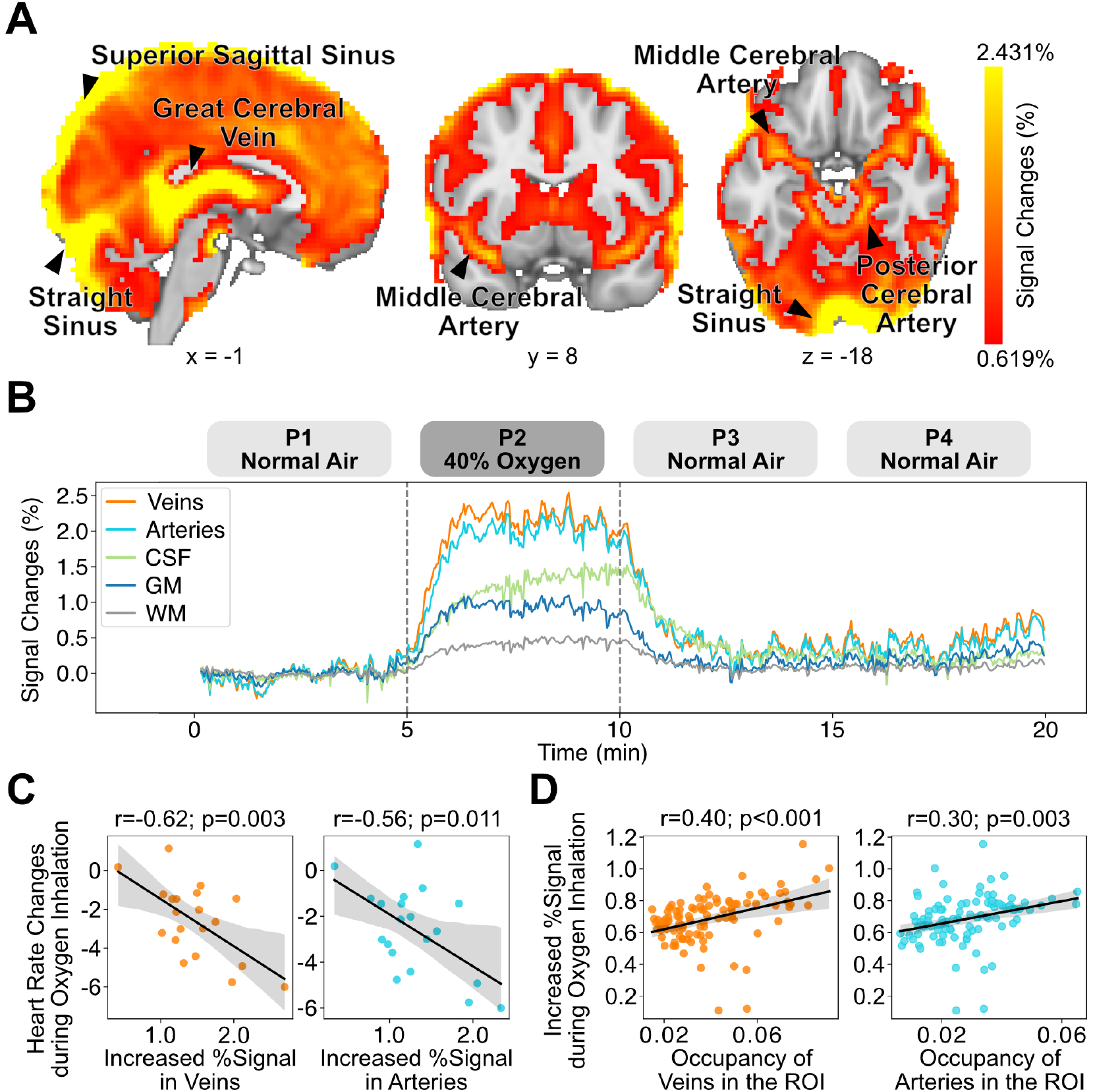
Analysis of BOLD signal changes induced by high-concentration oxygen inhalation on original data before the cleaning procedure. (A) Group-averaged oxygen-induced signal changes compared to baseline (P2–P1). The voxel-wise map of signal changes is thresholded at 50% of the highest value (0.619%) within the whole brain mask. The brightest color is 98% of highest value (2.413%). The greatest signal changes were observed in vascular regions, supporting the hypothesis that oxygen inhalation leads to global signal increases, primarily driven by changes in vascular areas. Major brain vessels are marked on the map. (B) Average signal time course across different brain tissues. Following oxygen inhalation at 5 min, signal increases were observed across the whole brain, with a drop at 10 min when supplemental oxygen was replaced with normal air. The highest signal changes occurred in veins and arteries, while white matter shows the smallest signal changes. (C) Participants with greater oxygen-induced reductions in heart rate exhibited larger signal changes in veins and arteries, suggesting that vessel oxygenation and heart rate reduction are linked to similar physiological processes. Each dot represents one participant. (D) Group-averaged signal changes in 100 neocortical GM ROIs as a function of the vascular occupancy within those regions. Each dot represents one ROI. Significant correlations indicated that GM regions with higher vascular occupancy (veins or arteries) exhibit greater signal changes. Vascular occupancy was determined by averaging the tissue probability within each GM region from the respective tissue probability maps.

To more precisely examine the dynamics of BOLD signals across different brain tissue, we segmented the cerebral cortex into gray matter (GM), white matter (WM), and cerebrospinal fluid (CSF). Additionally, we used publicly available cerebral vasculature templates (26) to define regions with highest veinous and arterial densities. We then extracted the time series signals from voxels associated with each tissue from individual runs (see Method). Bootstrapped mean signals across all the acquired data (Fig. 2B) showed that the signals in all tissue increased and saturated approximately 60 sec after administration of high-concentration oxygen inhalation (during P2). Following withdrawal of the high-concentration oxygen (during P3), the signals decreased and stabilized within about 60 sec. Additionally, individuals with greater reductions in heart rate following high-concentration oxygen inhalation exhibited higher signal changes in veins (Pearson’s correlation coefficient *r*_18_ = −0.62, p = 0.003, *BF*_10_ = 14.93; Cardiac data were available for 20 of the 22 participants) and arteries (Pearson’s correlation coefficient *r*_18_ = −0.56, p = 0.011, *BF*_10_ = 5.79; Cardiac data were available for 20 of the 22 participants, Fig. 2C). This outcome suggests that dissolved oxygen in the blood may serve as a common factor influencing both the observed reduction in heart rate and the magnitude of signal changes in veins and arteries.

To investigate BOLD signal changes beyond vascular contributions, we considered excluding voxels strongly associated with vessels and focusing solely on signal changes within GM. However, blood vessels are widely distributed in the brain, and collectively drain into venous sinuses. Consequently, signal increases observed in GM may still reflect contributions from these draining vessels, with regions containing higher vessel densities potentially exhibiting greater oxygen-induced signal changes. To evaluate this effect, we measured the average signal changes within each of 100 regions of interest (ROIs) in neocortical GM (27) and correlated them with the densities of veins and arteries within and around these ROIs. Vessel density was quantified as the average probability of veins and arteries based on respective tissue probability maps (see Methods). Our analysis revealed that brain regions with greater signal changes in GM were indeed associated with higher vessel density (Fig. 2D; Veins: Pearson’s correlation coefficient *r*_98_ = 0.40, p < 0.001, *BF*_10_ = 553.79; Arteries: Pearson’s correlation coefficient *r*_98_ = 0.30, p = 0.003, *BF*_10_ = 10.91). These findings underscore the importance of addressing vascular contributions when interpreting BOLD signal changes in GM. In the next section, we will explore methods to refine GM signal analysis by removing vascular contributions and reevaluating their impact.

### Eliminating Non-Neural Vascular Confounds in fMRI Signals Using ICA and Noise Classification

We aimed to isolate fMRI signal changes more specifically related to local neural activity by identifying and removing global vascular fluctuations derived from large-scale venous and arterial oxygenation changes associated with additional oxygen inhalation. To achieve this, we applied independent component analysis (ICA) to decompose the fMRI data into multiple components, each characterized by distinct spatial and temporal profiles, allowing us to identify and separate large-scale venous and arterial signals from those more closely linked to local neural activity. For instance, venous components typically exhibit spatial maps concentrated in the venous sinuses, particularly the sagittal sinus, and display a low-frequency temporal spectrum; whereas arterial components are spatially aligned with the great cerebral arteries, especially the middle cerebral branches pass through the insular cortex, and display a distinctive spectrum mixed by low and high frequencies (28).

To classify these components, we employed FMRIB’s ICA-based X-noiseifier (FIX), which computes spatial and temporal features for each component and uses a classifier to distinguish signal components from non-neurally driven ones (29, 30). This approach provides a reliable foundation to eliminate artifactual signals and improving the specificity of fMRI data analysis. We applied FIX with a pre-trained classifier (30) that was trained on hand-labeled datasets representing various artifacts, including global vascular artifacts from arteries and veins, as well as other artifacts from motion, WM, CSF, and other nonneural physiological sources (see details in Methods). This process identified approximately 44.85% of components as non-neurally driven one for individual fMRI runs (26.20 to 64.80%, SD = 8.48%; Supplementary Fig. 2A). We further verified that these non-neural components were more likely to be associated with non-GM regions (Supplementary Result 2, Supplementary Fig. 2B).

To clean the data, we regressed the time series of all identified non-neural components onto the voxel-wise signals in the original data. The resulting residuals were considered the cleaned data, with reduced influence from non-neural sources. The cleanup procedure effectively removed most signal fluctuations below 0.01 Hz (Supplementary Result 3, Supplementary Fig. 2C, D), which are typically attributed to extremely low-frequency drifts caused by non-neural source (31).

We then re-analyzed the oxygen-induced signal changes in the cleaned data and found that the overall magnitude of signal changes across the entire brain was substantially reduced (Fig. 3A color bar). For example, the median signal change decreased from 0.62% in the original data to just 0.02% after cleaning. This suggests that a large portion of the signal variation, likely driven by oxygen-induced global vascular responses, was effectively removed by the preprocessing procedure. This reduction was also apparent in the spatial distribution of the signal changes. Vascular regions such as the superior sagittal sinus and cerebral arteries no longer showed disproportionately large signal changes when compared to surrounding neocortical areas. However, vascular influences were not entirely eliminated. The great cerebral vein continued to exhibit the strongest signal changes, which may reflect localized venous oxygenation differences that were not fully addressed by the cleaning process. Consistent with the reduced signal in vascular regions, time-series extracted from different brain tissues showed that signal changes across various structures became less distinctive following cleaning (Fig. 3B). Notably, signal changes in veins and arteries were no longer associated with HR changes induced by high-concentration oxygen (Fig. 3C; veins: Pearson’s correlation coefficient *r*_18_ = −0.17, p = 0.467, *BF*_10_ = 0.35; arteries: Pearson’s correlation coefficient *r*_18_ = −0.11, p = 0.641, *BF*_10_ = 0.31), indicating that BOLD signal variations were dissociated from global vascular effects driven by high-concentration oxygen inhalation.

**Fig. 3.**
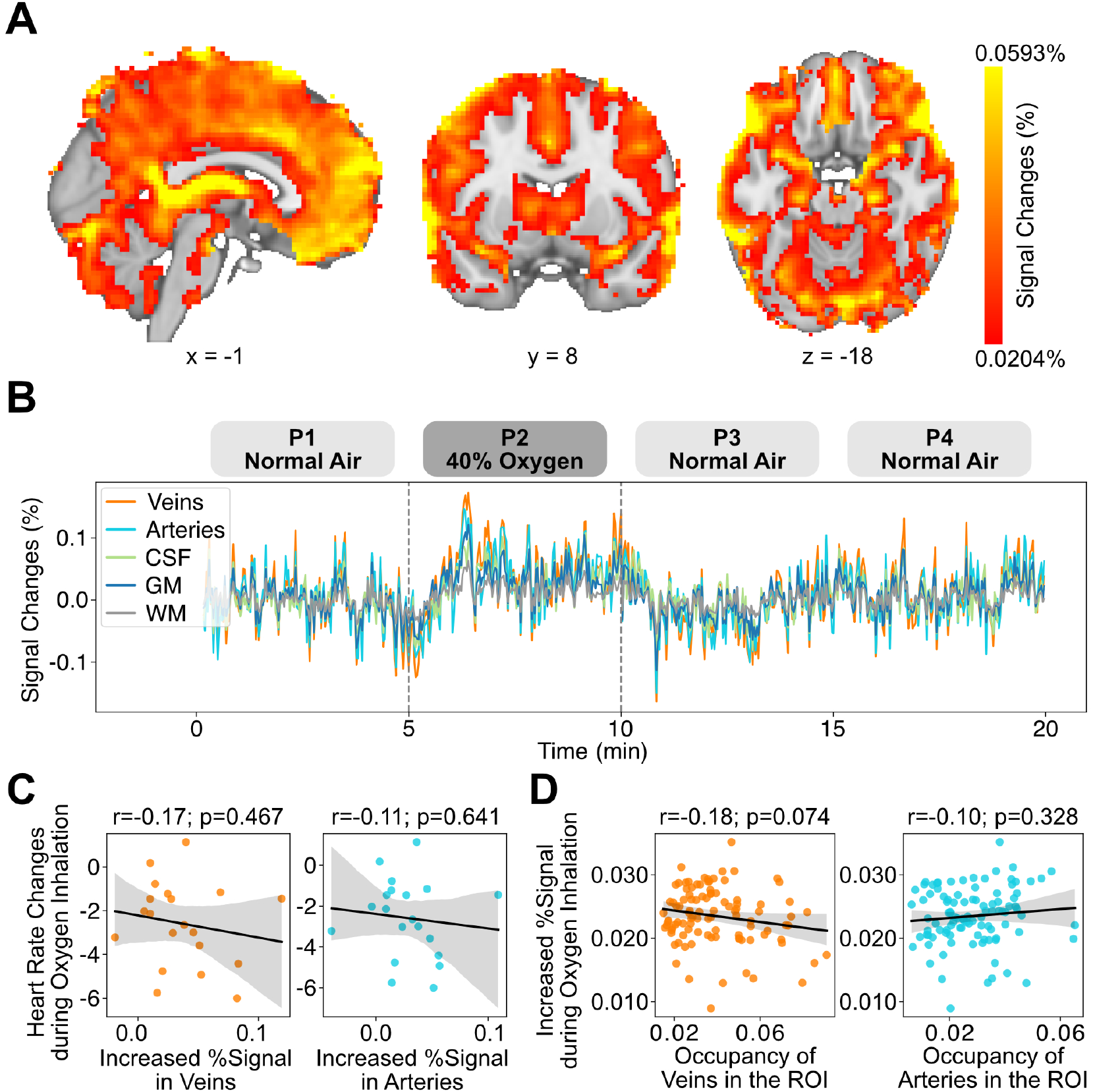
Analysis of BOLD signal changes induced by high-concentration oxygen inhalation on cleaned data after the cleaning procedure. (A) Group-averaged oxygen-induced signal changes compared to baseline (P2–P1). The voxelwise map of signal changes was thresholded at 50% of the highest value (0.0204%) within the whole brain mask. The brightest color is 98% of highest value (0.0593%). Compared to the original data in Figure 2, the magnitude of signal changes was significantly reduced after cleaning. (B) Average signal time course across different brain tissues. There was less distinction in signal changes between brain tissue types after the cleaning procedure. (C) With the cleaned data, heart rate no longer correlated with signal changes in veins and arteries, suggesting the cleaning procedure has removed this vascular influence. Each dot represents one participant. (D) The positive correlation between signal changes in GM regions and vascular occupancy no longer existed, suggesting that the data cleaning process effectively removed vascular contributions from GM regions. Each dot represents one ROI.

To avoid confounding effects from these non-cortical regions, subsequent analyses focused on GM. The oxygen-induced GM signal changes in the original data were strongly associated with vascular densities (Fig. 2D), underscoring the influence of vascular contributions. After applying the cleaning procedure, we expected that the residual signals in GM would primarily reflect neural activity, with minimal contribution from global vascular oxygenation induced by additional oxygen intake. To confirm this expectation, we re-examined the relationship between GM signal changes and vascular densities in the cleaned data. Crucially, the previously observed positive correlations in the original data were absent in the cleaned data (Fig. 3D, Veins: Pearson’s correlation coefficient *r*_98_ = −0.18, p = 0.074, *BF*_10_ = 0.60; Arteries: Pearson’s correlation coefficient *r*_98_ = 0.10, p = 0.328, *BF*_10_ = 0.20). This outcome supports the claim that the cleaning procedure effectively eliminated vascular contributions at least in GM, providing a foundation for analyzing non-vascular signal changes in GM regions. This critical step ensures that the residual GM signals have more dominant contribution from neural sources, allowing us to proceed with downstream analyses with greater confidence in the data’s integrity. In summary, we successfully employed ICA and noise classification techniques to identify and remove BOLD signal variations arising from vessels and other non-neural sources. By regressing out non-neural components, we obtained cleaned data that more closely represents localized neural activity, with reduced influence from systemic vascular oxygenation changes induced by high-concentration oxygen inhalation.

### “Brain Oxygen Sensitivity Topography” (BOST) — BOLD Dynamics During and After Oxygen Intake

Using the cleaned data, we investigated the spatial distribution and dynamics of BOLD signal changes induced by high-concentration oxygen inhalation. As shown in Fig. 3A, the spatial map reveals varying levels of signal increases across different cortical areas. To illustrate these differences more clearly, we averaged the signal changes within each of 100 neocortical GM ROIs, as defined by a publicly accessible parcellation method that identifies regions with homogeneous functional connectivity based on spatial proximity and assigns large-scale network labels according to global similarity (27). The results were then visualized on a cortical surface (Fig. 4A). This cortical surface representation highlights a heterogeneous response to high-concentration oxygen (P2–P1). We also compared the brain signals obtained during P3 and P4 to those occurring in P1 to examine the brain dynamics after oxygen inhalation; this process yielded BOST difference maps for both P3–P1 and P4–P1 (shown later). To clarify this procedure, we denote these BOST maps as BOST_*P*2−*P*1_, BOST_*P*3−*P*1_, and BOST_*P*4−*P*1_. Comparisons between adjacent experimental phases (e.g., P3–P2, P4–P3) were not conducted in this study, as we focused on contrasts relative to a common baseline (P1) to ensure consistent interpretability of temporal changes across phases.

**Fig. 4.**
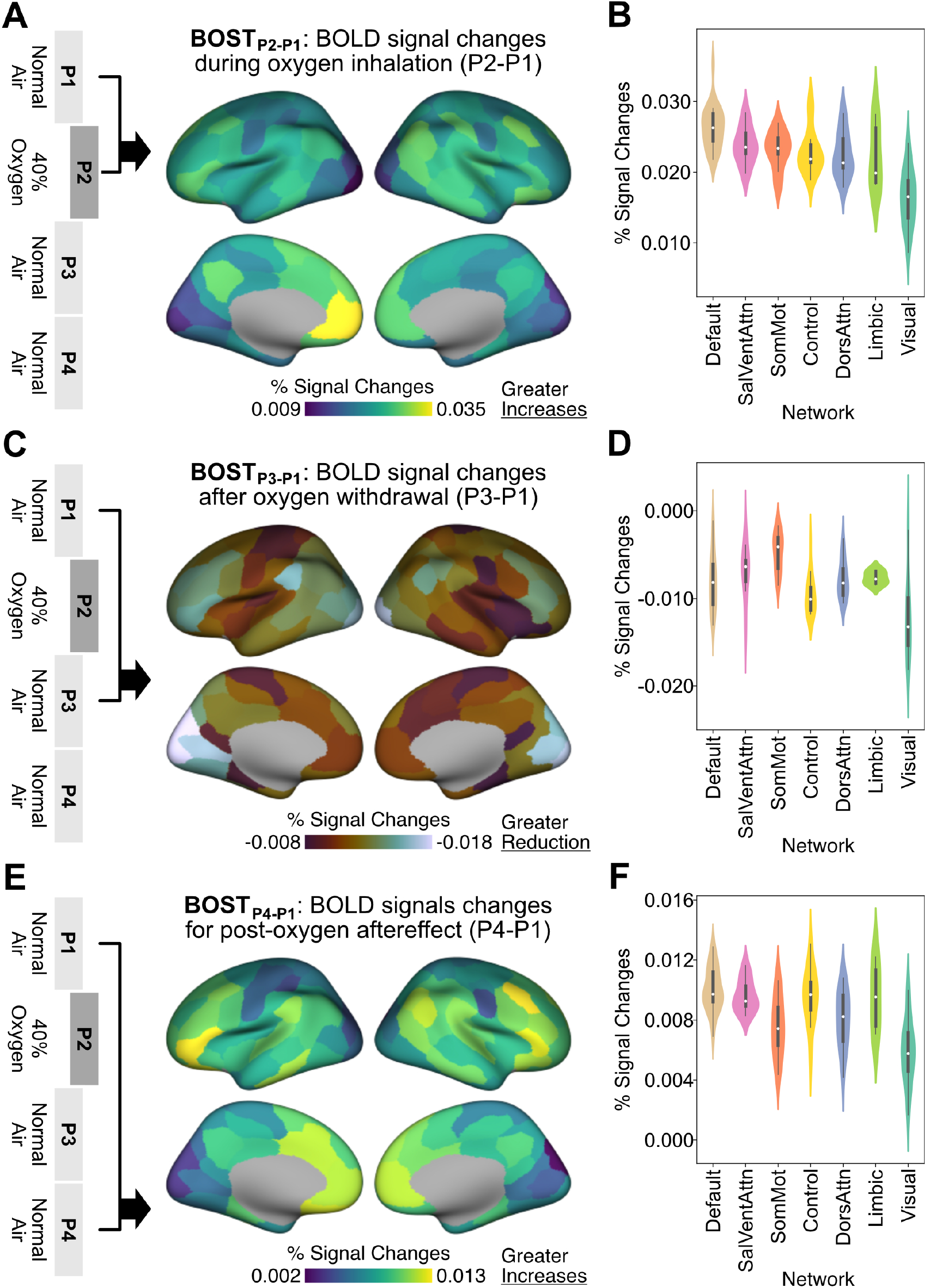
Brain Oxygen Sensitivity Topography (BOST) reveals spatial heterogeneity in BOLD signal changes across different phases. (A) Compared to the baseline phase (P1), GM regions showed varying degrees of signal change during oxygen inhalation (P2–P1), reflecting region-specific sensitivity to elevated oxygen levels. (B) When grouped by functional networks, these responses highlight spatial heterogeneity at the network level. Notably, the default mode network (DMN) showed a selectively heightened response to oxygen inhalation. (C) Following immediate oxygen withdrawal (P3–P1), BOLD signals decreased across nearly all regions, with the most pronounced reductions observed in the occipital cortex. (D) Network-level analysis revealed distinct patterns of signal reduction, suggesting differential dynamics in how networks responded to the removal of supplemental oxygen. (E) Surprisingly, BOLD signal levels began to rise again 5 min after oxygen withdrawal (P4–P1), even in the absence of continued oxygen delivery. (F) This rebound effect shows a spatial pattern partially resembling the original oxygen-induced activation, but with notably increased responses in the control and limbic networks, indicating potential delayed or lingering oxygen-related effects.

In BOST_*P*2−*P*1_, the most pronounced signal increases were observed in the medial prefrontal cortex (ROI centroid coordinates: x = −6, y = 46, z = 0, %signal changes = 0.035%). Lateral orbitofrontal cortex, precuneus, and inferior parietal lobule also showed higher amounts of increases compared to other cortical regions. These regions are predominantly associated with the DMN. When examining signal changes across networks (Fig. 4B), the mean value of signal change within the DMN had the highest signal changes compared to other networks. As a reference, we performed nonparametric permutation testing on voxelwise signal change maps from each individual to assess statistical significance (Supplementary Fig. 3, voxel-level family-wise error (FWE) correction at p < 0.05). The results of this analysis showed that most cortical regions, excluding the occipital cortex, exhibited significant modulation in response to high-concentration oxygen. Moreover, the statistical map also revealed that the bilateral hippocampus had significant effects of high-concentration oxygen (left hippocampus: cluster peak location: x = 26, y = −20 z = −16; cluster peak negative log(FWE p) = 3.40, cluster peak beta = 0.051%; right hippocampus: cluster peak location: x = 24, y = −22 z = −15.5; cluster peak negative log(FWE p) = 2.92, cluster peak beta = 0.066%). These findings underscore the widespread yet heterogeneous effects of inhaling high-concentration oxygen on brain activity, with diverse sensitivity patterns across GM regions.

We next investigated how these signals evolved after participants transitioned back to normal air. Specifically, we examined BOST_*P*3−*P*1_by comparing the signals during P3 to those during P1. Interestingly, after the withdrawal of high-concentration oxygen, the BOLD signals dropped to a level lower than the baseline (Supplementary Fig. 4A). Nonparametric permutation testing on voxel-wise difference map revealed statistically significant reductions only in a small portion of the left prefrontal cortex (Supplementary Fig. 4B, voxel-level FWE correction at p < 0.05; cluster peak location: x = −21, y = 41 z = 40; cluster peak negative log(FWE p) = 1.55, cluster peak beta = −0.02%). In BOST_*P*3−*P*1_(Fig. 4C), the most pronounced undershoots were observed in the occipital regions (ROI centroid coordinates: x = −6, y = −82, z = 26, %signal changes = −0.018%), corresponding to the visual network (Fig. 4D). In contrast, the weakest undershoots appeared in the somatomotor network. Notably, although the regional signal patterns between BOST_*P*2−*P*1_and BOST_*P*3−*P*1_were highly correlated (Supplementary Results, Supplementary Fig. 6A), the magnitude of the undershoots seen in P3–P1 did not mirror the increases observed for P2–P1. This asymmetry is also reflected in differences in network-level profiles between the two contrasts, which were not simply inverses of each other. P2–P1 showed smallest signal changes in the visual network and the largest in DMN (Fig. 4B). Whereas P3–P1 exhibited the most pronounced undershoot in the visual network and the weakest in the somatomotor network (Fig. 4D). These findings suggest that the return of BOLD signal to baseline after cessation of high-concentration oxygen does not occur in a symmetric manner across regions. Instead, BOLD signal dynamics relative to baseline show region-specific patterns, suggesting that the post-oxygen undershoot may involve physiological mechanisms distinct from those underlying the initial oxygen-induced increases.

We further examined BOST_*P*4−*P*1_by comparing the signals during P4 to those during P1 to determine whether the suppression observed in P3 persisted. Surprisingly, signals occurring during P4 were higher than those in P1 (Supplementary Fig. 5A), indicating another rise in the BOLD signal even after the withdrawal of high-concentration oxygen. Non-parametric permutation testing revealed robust increases, particularly in the frontal cortex (Supplementary Fig. 5B; cluster peak location peak location: x = −41, y = 15, z = 19; cluster peak negative log(FWE p) = 3.70, cluster peak beta = 0.0083%). BOST_*P*4−*P*1_(Fig. 4E) and its network summary (Fig. 4F), regions that exhibited higher oxygen-induced signal increases during P2 were more likely to show greater aftereffects during P4 (Supplementary Result 4, Supplementary Fig. 6B). However, notable differences in the spatial pattern and network profile compared to P2 were observed. During P4, while the medial prefrontal cortex continued to exhibit significant signal changes, the highest increases were found in the right lateral prefrontal cortex (ROI centroid coordinates: x = 44, y = 16, z = 44, %signal changes = 0.013%), a region associated with the control network. Notably, signal changes in the control network for P4–P1 were comparable to those in the default network (Default > Control for P4–P1: *t*_35_ = 0.88, p = 0.38, *BF*_10_ = 0.45). By contrast, for P2–P1, the control network exhibited significantly smaller signal changes compared to the default network (Default > Control for P2–P1: *t*_35_ = 3.13, p = 0.0035, *BF*_10_ = 11.30). These findings indicate that after the withdrawal of high-concentration oxygen, a secondary rise in BOLD signals occurs during P4, with increases most prominent in the control network. This suggests a delayed aftereffect that is partially related to oxygen-induced changes but also involves distinct spatial patterns that might be related to post-exposure-withdrawal mechanisms.

### Functional Decoding Predicts Potential Cognitive Functions That May be Affected by High-Concentration Oxy-gen

Inhalation and then withdrawal of high-oxygen yielded changes in BOLD signals across neocortical GM regions, characterized by a rise during P2, a subsequent fall during P3, and a secondary rise during P4. While the overall spatial patterns of signal changes were similar across these phases, notable distinctions emerged, thereby suggesting that high-concentration oxygen modulates different brain regions and differentially affected associated cognitive processes. To further understand how these oxygen-induced BOLD signal changes translated into its potential functional outcomes, we performed a functional decoding analysis. This analysis correlated voxel-wise signal change maps for each phase with meta-analytic maps of cognitive functions (32), enabling us to infer which cognitive domains were most closely associated with the observed activation patterns. This approach bridges the gap between signal changes to oxygen and their potential behavioral consequences, providing a basis for future studies on targeted oxygen-based cognitive modulation.

The top five cognitive functions whose meta-analytic maps show the strongest correlations with signal change patterns observed during high-concentration oxygen inhalation (BOST*_P_* _2−*P* 1_), its cessation (BOST*_P_* _3−*P* 1_), and its lasting effects (BOST*_P_* _4−*P* 1_) are shown in Fig. 5. Full lists of significantly correlated cognitive functions are available in Supplementary Fig. 7-9. The signal changes patterns observed in P2–P1 and P4–P1 were associated with meta-analytic maps corresponding to high-level cognitive functions such as social cognition, executive function, and language. Notably, the signal difference patterns identified in P4–P1 showed stronger associations with executive function and cognitive control compared to P2–P1. These stronger associations likely reflected greater engagement of the lateral prefrontal cortex observed in signal changes of P4–P1, suggesting a shift toward regulatory processes. By contrast, the physiology outcomes occurring during P3–P1 were linked to a more diverse array of cognitive associations, including attention, perception, semantics, and language. These differed from those cognitive associations found in P2–P1 and P4–P1. We observed the diverse patterns of signal changes suggest that oxygen interventions influence cognition in temporally- and spatially-specific ways. Functional decoding further provides a valuable framework for interpreting neural patterns in behavioral terms and offers insights into how oxygen administration may differentially influence specific cognitive domains across the phases of oxygen inhalation, withdrawal, and aftereffect.

**Fig. 5.**
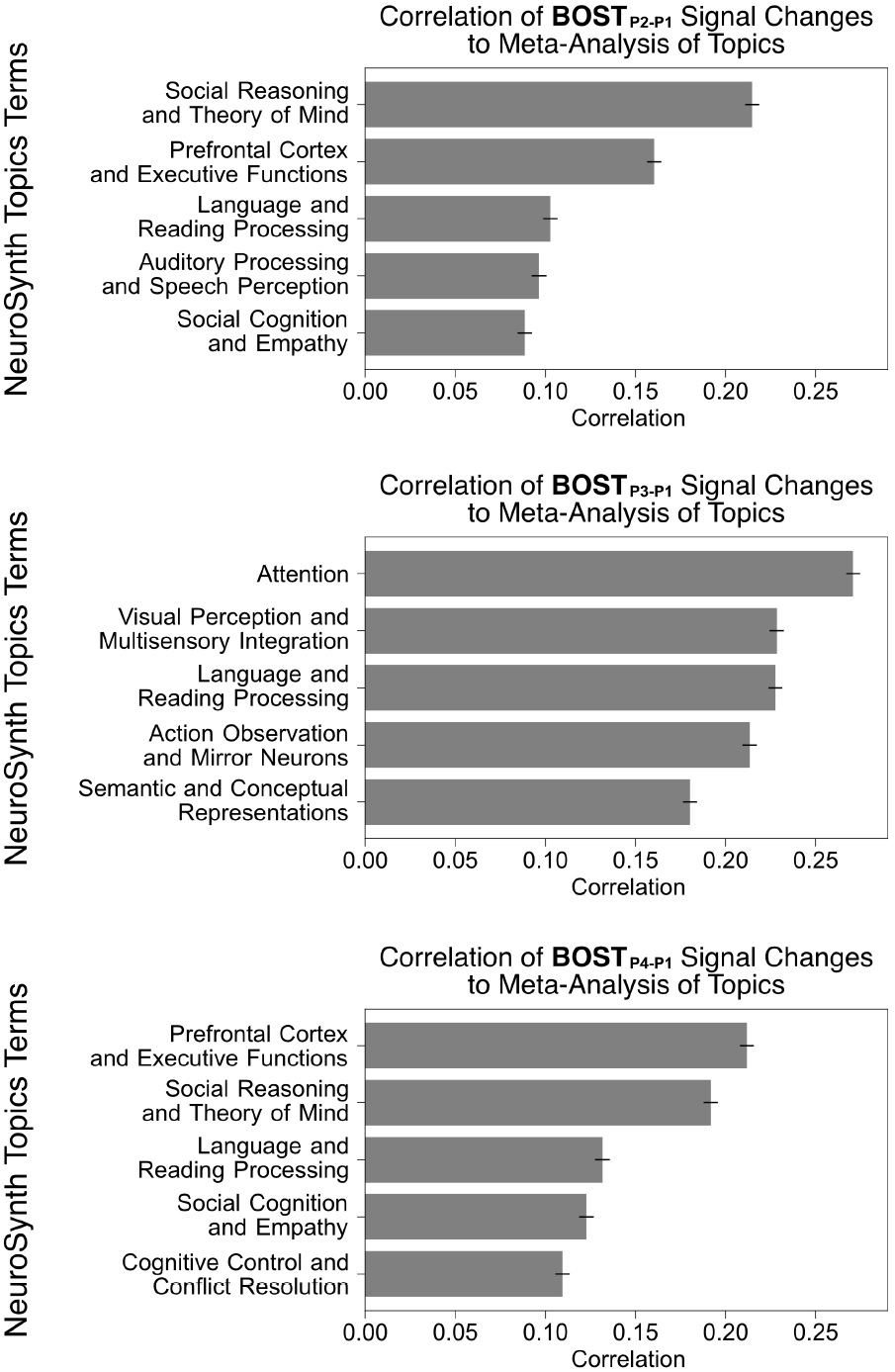
Functional decoding of spatial signal change patterns across different phases of high-concentration oxygen inhalation. Voxelwise BOLD signal change maps for each contrast, including oxygen inhalation (P2–P1), immediate oxygen withdrawal (P3–P1), and post-oxygen rebound (P4–P1), were functionally decoded using meta-analytic topic maps related to various cognitive functions. Each bar plot displays the top five cognitive topics whose spatial activation patterns showed the highest correlation with the observed BOLD signal changes. Bars represent correlation coefficients. The name of topic is a determined by several top terms under the topic.

### High-Concentration Oxygen Alters Between- and Within-Network Connectivity

High-concentration oxygen modulates BOLD signals in specific brain regions. Assuming that these enhanced BOLD signals reflect neural activity, the modified neural activation should then yield changes in the transmission of information across brain networks, leading to altering functional connectivity patterns. To investigate whether network reorganization occurred following changes in oxygen inhalation, we analyzed changes in functional connectivity across the different phases of oxygen inhalation levels. Time-series data from 100 neocortical GM regions were extracted separately for each the four phases and then used to construct pairwise functional connectivity matrices. These matrices were compared to P1 baseline functional connectivity, with statistical significance assessed using network-based statistics (33) to control the family-wise error rate (see details in Methods).

Our analysis revealed enhanced functional connectivity during high-concentration oxygen inhalation (P2–P1) and the post-oxygen aftereffect (P4–P1) (Fig. 6). However, no significant changes were observed for the immediate withdrawal phase (P3–P1). The enhanced connectivity during P2 was primarily associated with between-network connections, particularly between the DMN and other networks, such as the dorsal attention network, the salience/ventral attention network, and the somatomotor network. In contrast, functional connectivity changes during P4 were more extensive. In addition to enhanced between-network connections involving the DMN, within-network connections in the DMN itself were also significantly strengthened.

**Fig. 6.**
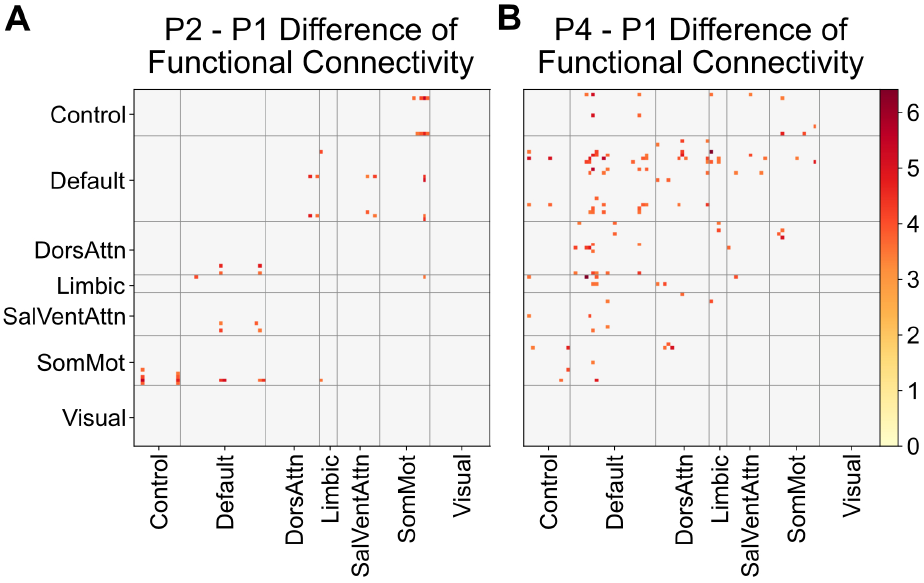
Changes in functional connectivity during and after high-concentration oxygen inhalation. Pairwise functional connectivity matrices for phases P2, P3, and P4 were computed and compared to baseline P1 connectivity. During oxygen inhalation (P2–P1), the default mode network (DMN) exhibited increased functional connectivity with multiple other networks. No significant connectivity changes were observed during the immediate withdrawal phase (P3–P1), and thus those results are not shown. In the post-oxygen phase (P4–P1), the DMN showed enhanced connectivity both within the network itself and with other large-scale brain networks.

These results indicate that high-concentration oxygen induces distinct changes in brain network connectivity across time points, with P2 characterized by increased inter-network coupling and P4 by both inter- and intra-network strengthening within the DMN. These findings underscore the dynamic and time-dependent effects of oxygen on functional brain network organization.

## Discussion

This study aimed to uncover the temporal dynamic and regional specificity of BOLD signal change during and after high-concentration oxygen intake. We hypothesized that excessive oxygen inhalation would induces global venous oxygenation, with localized neural activity appearing as superimposed changes. After removing vascular artifacts, the cleaned fMRI data revealed increased BOLD signals during oxygen inhalation, an undershoot after withdrawal, and reactivation even without continued oxygen. We mapped spatially distinct patterns in these dynamics, demonstrating the brain’s spatiotemporally heterogeneous responses to high-concentration oxygen, and suggest that different oxygen protocols have distinct effects on a wide range of cognitive functions.

### Oxygen Inhalation Shifts Energy Metabolism Especially in the DMN

Although the brain comprises only about 2% of the total body mass, it consumes nearly 20% of the body’s oxygen to support resting neural activity (34). To maintain energy homeostasis, the brain regulates two primary energy production pathways: mitochondrial oxidative phosphorylation and aerobic glycolysis. Oxygen is essential for oxidative phosphorylation, the brain’s main and most efficient pathway for generating ATP. By contrast, aerobic glycolysis does not rely on oxygen and provides energy more rapidly, though less efficiently. The balance between these two pathways shifts dynamically in response to fluctuations in oxygen and glucose availability (1). Empirical evidence shows that under high oxygen conditions, lactate levels decrease while overall glucose remains stable (35), indicating inhaling excessive oxygen can induce a shift from aerobic glycolysis toward oxidative metabolism. Additionally, elevated oxygen can lower carbon dioxide levels (36, 37), leading to vasoconstriction and a modest reduction (<10%) in cerebral blood flow (36, 38). This reduction may restrict glucose delivery, further favoring oxidative metabolism, which yields more energy per glucose molecule. Thus, high-concentration oxygen inhalation triggers a shift toward oxidative processing to produce an excess of energy. This surplus of energy supports the restoration of ion pumps and promotes sustained neuronal activity, which might contribute to the increases in neural activity and raised blood oxygenation levels, ultimately leading to the heightened BOLD signals observed in our study.

The DMN appears particularly sensitive to these aforementioned metabolic shifts. Our findings showed that both oxygen inhalation and the subsequent reactivation phase preferentially modulated BOLD activity within the DMN. This observation is consistent with prior studies reporting that hypoxia selectively alters cerebral blood flow in DMN regions (39) and that increased glucose availability can suppress DMN activity and connectivity (40). Prior research also suggests the DMN consumes more glucose than required for oxidation alone (41, 42), possibly to support its complex connectivity and cognitive functions, although other work challenges this view (43, 44). Taken together, these findings suggest that the heightened sensitivity to high-concentration oxygen inhalation in our study may stem from the unique metabolic characteristics of DMN.

### Variant Regulators Might Shape Rapid and Prolonged Modulation Leading to Distinct Temporal Dynamics

We observed a characteristic pattern in the BOLD signal following oxygen inhalation. The signal increased during inhalation, decreased below baseline immediately after withdrawal, and then rose again even when oxygen returned to regular levels. These changes likely reflect distinct immediate and sustained biological responses that regulate glucose and oxygen metabolism to maintain energy balance. For example, oxygen withdrawal induces a relative hypoxic state, activating hypoxia-inducible factor-1 (HIF-1), which shifts cellular metabolism toward nonoxidative glycolysis to maintain energy production under reduced oxygen conditions (45). This metabolic adjustment likely explains the undershoot in the BOLD signal observed immediately after oxygen withdrawal. Furthermore, high oxygen levels can reduce the availability of nitric oxide (NO), impairing its role in inhibiting mitochondrial electron transport chain activity (46–49). Notably, changes in nitric oxide levels can persist for at least 15 min after oxygen administration ends (50), suggesting that the metabolic effects of oxygen extend beyond the inhalation phase. Nevertheless, AMP-activated protein kinase (AMPK) senses changes in the AMP/ATP ratio and upregulates glucose transporters (GLUTs) to facilitate energy production (51–53). This regulation involves transcriptional control, such as the upregulation of the GLUT4 gene, promoting long-term metabolic adaptations (54, 55). These processes contribute to longer term metabolic adjustments following oxygen exposure. Together, these mechanisms provide both rapid responses and prolonged effects on brain metabolism and might explain the observed dynamic BOLD signal changes that occur after oxygen inhalation and withdrawal.

### Distinct Temporal and Regional Brain Responses May Facilitate Different Cognitive Functions

While the DMN consistently shows strong BOLD signal increases during both high-concentration oxygen inhalation and withdrawal, we observed a disproportionately higher activation in the lateral prefrontal cortex specifically after withdrawal. This suggests that the effects of high-concentration oxygen are both time-dependent and region-specific, potentially supporting distinct neural processes across different cognitive domains. Previous studies have shown that variations in the timing and duration of high-concentration oxygen exposure can lead to different cognitive improvements. For instance, short bursts (30 sec to 1 min) of high-concentration oxygen enhance short-term memory and performance on cognitively demanding tasks (9, 24). Continuous oxygen delivery during verbal learning or working memory tasks has also been shown to improve behavioral performance (8, 11). Moreover, prolonged high-concentration oxygen exposure (>1 hour) over multiple consecutive days has been associated with improvements in working memory, short-term memory, and information processing speed (56, 57).

Although several studies have reported cognitive improvements following high-concentration oxygen administration, there is still no clear consensus on the optimal timing, duration, or method of delivery. Differences in experimental designs and task types have led to inconsistent findings, making it difficult to draw firm conclusions about how the dynamics of high-concentration oxygen exposure influence specific cognitive processes. To explore this issue further, we applied functional decoding to identify which cognitive functions are most closely associated with BOLD signal increases observed at different phases of high-concentration oxygen inhalation and withdrawal. The results showed that the meta-analytic maps for social cognition, executive control, and language closely aligned with regional signal increases observed in both the P2–P1 and P4–P1 contrasts. This suggests that oxygen supplement induce additional neural activation specifically in the regions associated with these cognitive functions, potentially enhancing related behavioral outcomes. These cognitive enhancements are expected to be more pronounced for functions with stronger associations compared to those with weaker associations, such as emotion and decision-making (see Supplementary Figures 7–9 for the full ranking). Notably, the functional decoding results revealed time-specific patterns. Withdrawing from short bursts oxygen may be especially effective for enhancing cognitive control. Overall, functional decoding offers a framework for future studies to develop hypotheses and predictions regarding the behavioral effect of high-concentration oxygen, especially in terms of which cognitive functions are most affected and the timing of their enhancement.

Additionally, we observed distinct patterns of functional connectivity reorganization during high-concentration oxygen inhalation and the post-oxygen aftereffect, reflecting dynamic modulation of resting-state network architecture. High-concentration oxygen inhalation primarily enhanced between-network connectivity, promoting integration of information across functionally distinct systems. Such integration may facilitate coordinated processing of complex cognitive tasks that draw on multiple cognitive domains, consistent with prior findings linking between-network connectivity to general cognitive ability (58). In contrast, the post-oxygen aftereffect further strengthened within-network connectivity in the DMN. Such within-network connections may reflects increased temporal synchrony among DMN regions, promoting the integration of internally generated information within this network while reducing interference from other networks. This specialization may enhance the functional role of the DMN in various internally-driven processes, such as introspection (59), theory of mind (60), and creative thinking (61). These results suggest that oxygen levels can actively reshape the brain’s resting-state network organization even in the absence of external tasks or clinical conditions, offering new insights into how physiological factors shape intrinsic neural communication.

### Limitations

To isolate BOLD signal changes related to neural activity, we analyzed GM signals from the residuals after regressing out non-neural components. This approach assumes that hyperoxia-induced venous signals are linear (without nonlinear interactions between different non-neural sources), symmetric (proportional increases and decreases), and time-invariant (without dramatic delays or latency) fluctuations. However, our analysis of the cleaned data revealed the persistence of large signal changes in the great cerebral vein suggests that some venous regions violate these assumptions, likely due to the specifically localized oxygenation in nearby deep brain structures. While our cleaning procedure reduced much of the non-neural noise in GM, it could not fully eliminate venous confounds in the whole brain. Notably, venous bias remains a common challenge in fMRI studies (62–65), and improved methods for removing such artifacts are essential for advancing neuroimaging research for oxygen inhalation.

While BOLD fMRI presents challenges under oxygen inhalation, our findings reveal temporal dynamics in region-specific enhancements and functional network changes, offering valuable spatial insights that complement previous electrophysiological studies, which lack spatial resolution. Although each neurophysiological modality has its own constraints, they often provide complementary perspectives. Thus, future studies should consider multimodal approaches to obtain more comprehensive and converging evidence to better address the complexities of non-neural contributions. For example, combining fMRI with simultaneous EEG could help disentangle vascular and neuronal effects by offering high temporal resolution of neural activity alongside spatially detailed hemodynamic data (66). Additionally, incorporating Arterial Spin Tag Labeling could provide valuable insights into cerebral blood flow to more accurately identify vascular contributions in the cortical brain regions in each individual (67, 68). Moreover, as previously noted, glucose metabolism may play a critical role under hyperoxic conditions. Thus, incorporating positron emission tomography (PET) imaging provide more direct measurement of regional glucose and oxygen consumption following high-concentration oxygen inhalation, further clarifying the underlying physiological mechanisms (69).

In summary, our findings underscore the temporal dynamics and region-specific effects of oxygen inhalation on BOLD signals, offering new insights into the temporal and spatial dimensions of brain activity. Isolating neural signals from vascular influences remains challenging, and multimodal approaches are recommended in future studies for more accurate and direct assessments of the many factors involved. Overall, examining the effects of oxygen inhalation modulated BOLD signals promises to deepen our understanding of the interplay between neural and vascular responses, and clarify metabolic mechanisms driving oxygen-induced cognitive modulation.

## Method

### Participants and experimental procedure

We recruited participants through Jikken-baito, an online platform that distributes experiment advertisements across multiple social media channels to connect researchers with potential participants. Participants were screened for eligibility to undergo MRI and excluded if presented psychiatric, neurological, cardiac and respiratory disorders. All participants provided informed consent, which was approved by the Research Ethics Committee of the University of Tokyo. Each participant completed two functional imaging runs while alternating between normobaric high-concentration oxygen and normal air inhalation. Oxygen was supplied through a medical-grade oxygen concentrator (DAIKIN LiteTEC-5B), while normal air was delivered via an air pump. The run began with 5 min of normal air inhalation, followed by 5 min of oxygen delivery, and concluding with 10 min of normal air. The airflow protocol was identical across participants, who were blinded to the timing of air and oxygen delivery. Participants were given a break of at least 1 min after the first scanning run and began the second run when they were ready.

### Data acquisition

The MRI data were collected with a 3T Siemens MAGNETOM Prisma MRI scanner using a 32-channel head coil. Structural T1-weighted images were acquired using an MPRAGE sequence with the following parameters: 192 sagittal slices, FoV = 240 × 256 mm, matrix = 240 × 256, TR = 2400 ms, TE = 2.15 ms, flip angle = 8°, and slice thickness = 1 mm. T2-weighted structural images were acquired with 192 sagittal slices, FoV = 240 × 256 mm, matrix = 240 × 256, TR = 3200 ms, TE = 564 ms, flip angle = 120°, and slice thickness = 1 mm. Functional images were collected using a multiband echo-planar imaging (EPI) protocol with the following parameters: FoV = 195 × 195 mm, matrix = 78 × 78, TR = 2000 ms, TE = 30 ms, flip angle = 74°, 54 slices covering the whole brain, slice thickness = 2.5 mm, and a multiband factor of 2. Each participant completed two functional scanning runs, with each run consisting of 1200 EPI volumes. To correct for field inhomogeneity, two spin-echo field maps were acquired, one with an anterior-to-posterior (A → P) phase encoding direction and the other with a posterior-to-anterior (P → A) direction.

### Physiological recordings

Cardiac and respiratory signals were recorded using the Siemens built-in physiology monitoring unit (PMU) and an abdominal breathing belt, sampled at 200 Hz and 50 Hz, respectively. The analysis of these signals was performed using `neurokit2`, which included preprocessing, peak detection, and the calculation of key metrics such as heart rate (HR), heart rate variability (HRV), respiratory rate (RR), respiratory volume per time (RVT), and respiratory rate variability (RRV). We used a two-way repeated measures ANOVA to examine differences in the metrics across four phases and two restingstate runs. Post-hoc comparisons with Tukey adjustment were performed using the `emmeans` package in R.

### fMRI data preprocessing

No participants were excluded due to excessive head motion; we set the exclusion threshold at a mean framewise displacement exceeding 0.9 mm or more than 20% of functional imaging volumes with framewise displacement greater than 0.9 mm (70). The fMRI data underwent minimal preprocessing using `fMRIPrep`, including transformations into MNI space. However, our subsequent cleaning procedures were performed on the preprocessed images in native space. The resulting cleaned data were then transformed into MNI space. These functional images were further processed with 5 mm spatial smoothing and a mild high-pass temporal filter (>2000 s) to remove linear drift. We then applied FSL’s MELODIC for independent component analysis (ICA) and used FIX-ICA with the pre-trained “Standard.RData” dataset to identify venous and other non-neural artifacts components. Before removing artifactual components, motion parameters were regressed out, and this intermediate dataset was considered as “original data”, which presumably retained significant vascular contributions. To obtain “cleaned data”, we then regressed out the identified artifactual components using FIX-ICA’s aggressive option. Both the original and cleaned datasets were transformed to MNI space and scaled to represent percent signal change, using the first 5 min (during inhalation of normal air) as the baseline.

### BOLD signal change analysis

We analyzed signal changes using `nilearn`. The average BOLD signal for four 5-min phases were estimated using a general linear model (GLM) with four discrete boxcar functions, one for each phase, and an autoregressive model (AR[1]) to account for autocorrelated noise. We made three contrasts (P2–P1, P3–P1, and P4–P1) across two runs based on the estimated boxcar functions to assess BOLD signal changes between phases. These contrasts were then entered into a second-level GLM to estimate the group-average differences between phases. To determine statistical significance, we performed 10,000 non-parametric permutations using a one-sample t-test with a sign-flipping strategy to generate the null distribution. The resulting statistical maps were thresholded using voxel-level family-wise error (FWE) correction at p < 0.05.

We further extracted representative time courses from different brain tissues. Voxel selection was guided by tissue probability maps for GM, WM and CSF (71) as well as artery and vein (26). Due to substantial interindividual variability in vascular anatomy, and generally lower probability values in artery and vein maps, the same selection criteria used for GM, WM, and CSF could not be uniformly applied across all tissue types. Instead, we selected a fixed number of voxels with the highest probability values: 10,000 for GM, 10,000 for WM, and 500 each for CSF, arteries, and veins. The numbers of voxels were arbitrarily determined but chosen to approximate the relative proportions of each tissue type in the brain while ensuring spatial specificity. The selections were visually inspected to ensure that the location of selected voxels plausibly represented each tissue. These arbitrary numbers resulted in tissue probabilities thresholded at 0.909 for grey matter, 0.989 for white matter, 0.970 for CSF, 0.777 for veins, and 0.676 for arteries. Time courses were extracted from the selected voxels and averaged to capture the temporal dynamics of each tissue type for each run. To further characterize BOLD signal changes across subjects, individual time courses were averaged using a bootstrapping method, providing a robust estimate of tissue-specific BOLD fluctuations.

To further explore the regional variability of oxygen’s effects on the brain, we extracted BOLD signal change estimates using Schaefer’s 100-region gray matter parcellation (27). This parcellation procedure assigned each region of interest (ROI) to a functional network, enabling us to determine the network affiliations of the affected ROIs. While the method supports multiple spatial resolutions (e.g., 100 to 1000 ROIs), we selected the 100-region version to balance spatial specificity with interpretability. Specifically, the 100-region resolution provides sufficient anatomical detail while maintaining a manageable number of ROIs, which facilitates clearer visualization and interpretation of results. We hypothesized that vessel densities in the gray matter might contribute to oxygen-induced BOLD signal changes. To investigate this relationship, we extracted artery and vein probability values from their respective tissue probability maps (as described in the previous paragraph). These probability values were then averaged across voxels within each region of interest (ROI) to obtain a regional measure of vascular density. Finally, we correlated these vascular density estimates with group-averaged BOLD signal changes from each ROI to assess whether regions with higher vessel densities exhibited greater oxygen-induced signal fluctuations.

### Functional decoding

We retrieved the Neurosynth dataset version 7 with 50 topics derived from topic modeling methodology and conducted a meta-analysis with `nimare` to generate functional maps for each topic. We then correlated BOLD signal change maps with these functional maps. The correlation coefficient represents the similarity between the functional mapping of a specific topic and the oxygen-induced BOLD signal changes. Statistical significance of correlation was corrected using Bonferroni correction. We asked ChatGPT to generate descriptive labels based on the most strongly associated terms. The terms for each topic were manually reviewed terms and specifically marked the topic that have merely association with cognitive function. In the main text, only topics with cognitive function were shown. The full lists of significant correlations are shown in supplementary results.

### Functional Connectivity Analysis

Functional connectivity was calculated by extracting time courses from each of the 100 GM ROIs and segmented into four time phases. To minimize confounding effects from immediate changes following airflow switches, we focused on the final 4 min (120 TRs) of each phase. The time courses from each phase were further applied bandpass filter (low pass = 0.083 Hz, high pass = 0.0078 Hz) and standardized to z-value. Pairwise Pearson correlations were then computed among the 100 ROIs to assess connectivity for each phase in each run. The connectivity matrix from two run were averaged to represent the functional connectivity for each individual and submitted to paired t-test comparison between phases. To identify significant differences across phases while controlling for the false discovery rate, we applied non-parametric network-based statistical analysis with 10000 permutation, the family-wise error rate were controlled by the clusters in connected graph components with the statistic thresholded at 3.433, equivalent to uncorrected p-value 0.001 (33).

## Data availability

The data that support the findings of this study are available from the corresponding authors upon reasonable request.

## Code availability

Code for performing the analyses and generating the visualizations is publicly available on GitHub at https://github.com/yushiangsu/highoxygenfMRI_2025.

## Author contributions

Y.S. collected the brain imaging data, conducted the experiments and data acquisition, performed data analysis and results visualization, and wrote the initial draft of the manuscript; C.W. conceived and designed the study, set up the equipment and experimental procedures, and conducted the experiments and data acquisition; J.G. provided methodological consultation, contributed to the interpretation of results, and reviewed the manuscript; J.S. provided methodological consultation, contributed to the interpretation of results, and reviewed the manuscript; Z.C. conceived and designed the study, set up the equipment, supervised the project, and provided funding; All authors approved the final version of the manuscript.

## ACKNOWLEDGEMENTS

Y.S., C.W., Z.C. discloses support for the research of this work from IRCN-Daikin SCP. Y.S. discloses support for the research of this work from JSPS KAKENHI Grant-in-Aid for Research Activity Start-up [grant number 24K22796]. Z.C. discloses support for the research of this work from World Premier International Research Center Initiative (WPI), MEXT, Japan.

## Supplementary Result 1: Effects of High-concentration Oxygen on Cardiac and Respiratory Responses

Due to hardware issues, we excluded data from two participants with invalid cardiac signals and from another four participants with invalid respiratory signals. The remaining valid data were preprocessed to compute multiple common metrics, including heart rate (HR), heart rate variability (HRV), respiratory rate (RR), respiratory volume per time (RVT), and respiratory rate variability (RRV). Repeated-measures ANOVA was used to analyze metric changes across phases (P1-P4), resting-state runs (i.e. the first or second 20-min run) were included as a secondary factor to control for potential time-related drift over the course of the experiment. We primarily focused on the phase effect to assess the impact of high concentration oxygen on physiological responses, and we further applied post-hoc pairwise comparison to examine if the oxygen inhalation has significant changes that is different from the other phases.

Among the five metrics derived from the cardiac and respiratory signals, HR is the only physiological metric showing a significant main effect of phases (*F*_3,57_ = 10.65, p < 0.001, *η*^2^ = 0.36) without a significant effect of resting-state runs (*F*_1,19_ = 1.39, p = 0.25, *η*^2^ = 0.07), indicating notable changes between phases (Fig. 1C).

RVT (Supplementary Fig. 1) exhibited significant main effects of phases (*F*_3,51_ = 4.59, p = 0.006, *η*^2^ = 0.21) and resting-state runs (*F*_1,17_ = 9.74, p = 0.006, *η*^2^ = 0.36). Further post-hoc analysis indicated the last phase have significant lower RVT than the first phase (P4 > P1: *t*_51_ = −3.50, p = 0.005, *η*^2^ = 0.19), and RVT in the second resting-state run is lower than first run (*t*_17_ = 3.12, p = 0.006, *η*^2^ = 0.36). These results suggest that RVT declined over time but was not specifically modulated by high-concentration oxygen. No significant phase effects were observed for other physiological metrics (Supplementary Fig. 1), HRV: *F*_3,57_ = 0.25, p = 0.86, *η*^2^ = 0.01; RR: *F*_3,51_ = 0.95, p = 0.424, *η*^2^ = 0.05; RRV: *F*_3,51_ = 1.96, p = 0.13, *η*^2^ = 0.10). No respiratory metrics showed a specific association with oxygen inhalation. This lack of significant respiratory changes is consistent with previous findings on normobaric high-concentration oxygen (10, 23), However, the study using hyperbaric oxygen have reported a decrease in respiratory rate (22). Further analysis is required to clarify the physiological mechanisms underlying these differences, particularly how oxygen concentration and pressure interact to modulate respiratory function.

## Supplementary Result 2: Evaluation of spatial characteristic of non-neural components for cleanup procedure

To evaluate the accuracy of this classification for identifying non-neuronal effect on our data, we examined the spatial features of both signal and non-neural components. We did this by thresholding component weights (greater than 3 or less than −3) to generate topographical maps and then assessing the proportions of different brain structures represented in those maps. We assessed the differences between non-neural and signal components using separate linear mixed-effect models for each brain structure. In each model, a binary indicator distinguishing non-neural from signal components was included as a fixed effect predictor, while subject was modeled as a random effect. The dependent variable was the probability of a given brain structure for each component. We used non-parametric bootstrapping with 10,000 simulations to estimate the beta coefficients for the binary indicator variables for each model. These coefficients reflect the difference in structure probabilities between non-neural and signal components. Confidence intervals were adjusted for multiple comparisons using the Bonferroni method. We found that signal components were predominantly associated with GM and WM, while non-neural components had significantly higher probabilities of being associated with veins, arteries, and CSF (Supplementary Fig. 2B; non-neural > signal components in probabilities of veins: estimated difference = 0.71%, 95% CI = [0.65, 0.76]; arteries: estimated difference = 0.63%, 95% CI = [0.57, 0.68]; GM: estimated difference = −5.86%, 95% CI = [-6.13, −5.57]; WM: estimated difference = −0.63%, 95% CI = [-0.98, −0.26]; CSF: estimated difference = 1.56%, 95% CI = [1.39, 1.73]). These results highlight the effectiveness of FIX in distinguishing vascular and other non-neuronal contributions from genuine neuronal signals, especially for veins, arteries, and CSF, thereby strengthening the validity of subsequent analyses using the cleaned data.

## Supplementary Result 3: Examining Temporal Signal Features Before and After Denoising

To assess whether the data cleaning procedure effectively preserved meaningful temporal features of the BOLD signal within the range of 0.01 to 0.1 Hz (72–74), we calculated temporal spectrums for each 5-min phase to examine the temporal characteristics of the original and cleaned datasets (Supplementary Fig. 2C). In the original data, there was a noticeably elevated power of extremely low-frequency signals (< 0.01 Hz) across all four phases. Notably, the oxygen inhalation phase P2 exhibited significantly higher power in this range compared to the baseline phase P1 (Supplementary Fig. 2D; *t*_21_ = 2.71, p = 0.013). Such fluctuations are often associated with non-neuronal sources, including slow head motion, scanner drift, respiratory effects, and changes in end-tidal carbon dioxide (31, 73, 75), and may be further amplified by oxygen-related physiological changes.

Importantly, the cleaning procedure effectively removed most of the fluctuations below 0.01 Hz, while preserving power within the functionally relevant range of 0.01-0.1Hz (Supplementary Fig. 2C). After cleanup, the previously observed enhancement in extremely low-frequency power during oxygen inhalation was not observed in the cleaned data (Supplementary Fig. 2D; *t*_21_ = 1.61, p = 0.12), indicating that oxygen-related noise was successfully removed. These results suggest that the cleaning procedure effectively attenuated non-neuronal signal fluctuations, particularly those in the extreme low-frequency range, while maintaining the temporal fidelity of neuronal signal fluctuations. This ensures the integrity of following analyses with the cleaned dataset.

## Supplementary Result 4: Distinct and Shared Regional Dynamics Across Oxygen Inhalation Phases

To examine the similarities in regional BOLD responses during different phases of oxygen inhalation and withdrawal, we computed correlations between group-averaged regional responses from BOST_*p*2−*p*1_, BOST_*p*3−*P*1_, and BOST_*p*4−*p*1_(Supplementary Figure 6). The strongest correlation was observed between signal changes patterns in P2–P1 and P4–P1 (Pearson’s correlation coefficient *r*_98_ = 0.811, p < 0.001), followed by a moderate correlation between signal changes patterns in P2–P1 and P3–P1 (Pearson’s *r*_98_ = 0.385, p < 0.001). The weakest, though still positive, marginal significant correlation was found between signal changes patterns in P3–P1 and P4–P1 (Pearson’s correlation coefficient *r*_98_ = 0.195, p = 0.052). Interestingly, despite P2 and P4 being the most temporally separated, their regional response patterns were the most similar. This suggests that the magnitude of regional signal changes may follow a stable temporal trend, where regions with greater signal increases during oxygen inhalation (P2) tend to show smaller undershoots during withdrawal (P3) and stronger reactivations during the post-oxygen phase (P4). However, as noted in the main text, this trend is not consistent across all brain regions. For example, the somatosensory cortex exhibited relatively low signal increases during P2 but did not show pronounced undershoots in P3 (Fig. 4A, 4C). Similarly, the lateral prefrontal cortex displayed a relatively stronger signal increase in the later P4 phase compared to P2 (Fig. 4A, 4E). These findings imply that, beyond a general trend driven by a global effect, region-specific and time-specific mechanisms may also shape the dynamics of oxygen-induced BOLD signal changes across different phases.

**Supplementary Figure 1.**
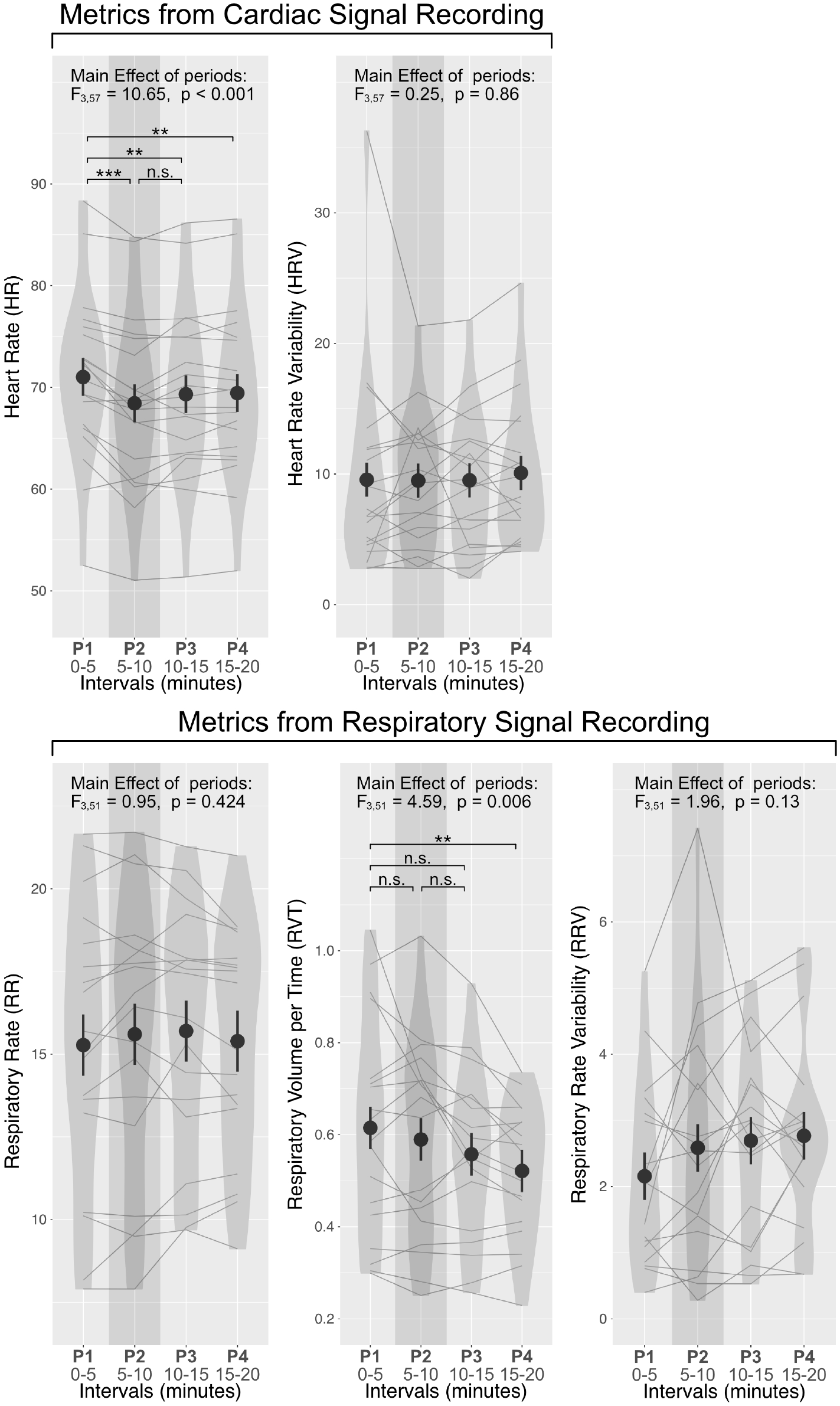
Effects of high-concentration oxygen on cardiac and respiratory responses. Cardiac and respiratory signals were preprocessed to compute indices for heart rate (HR), heart rate variability (HRV), respiratory rate (RR), respiratory volume per time (RVT), and respiratory rate variability (RRV). The average value of each metric for each phase was analyzed using repeated-measures ANOVA, with statistical results reported at the top. Among these metrics, only HR and RVT showed significant effects across phases. Post-hoc pairwise comparisons were conducted to determine whether these effects were specifically associated with oxygen inhalation during P2 (shaded area). HR decreased significantly when participants inhaled oxygen, while RVT exhibited a linear drift, with significant differences observed only between the first (P1) and last (P4) phases. Dots and error bars represent the mean and standard error of metric at each phase, while each line represents individual participants’ trajectories over time.

**Supplementary Figure 2.**
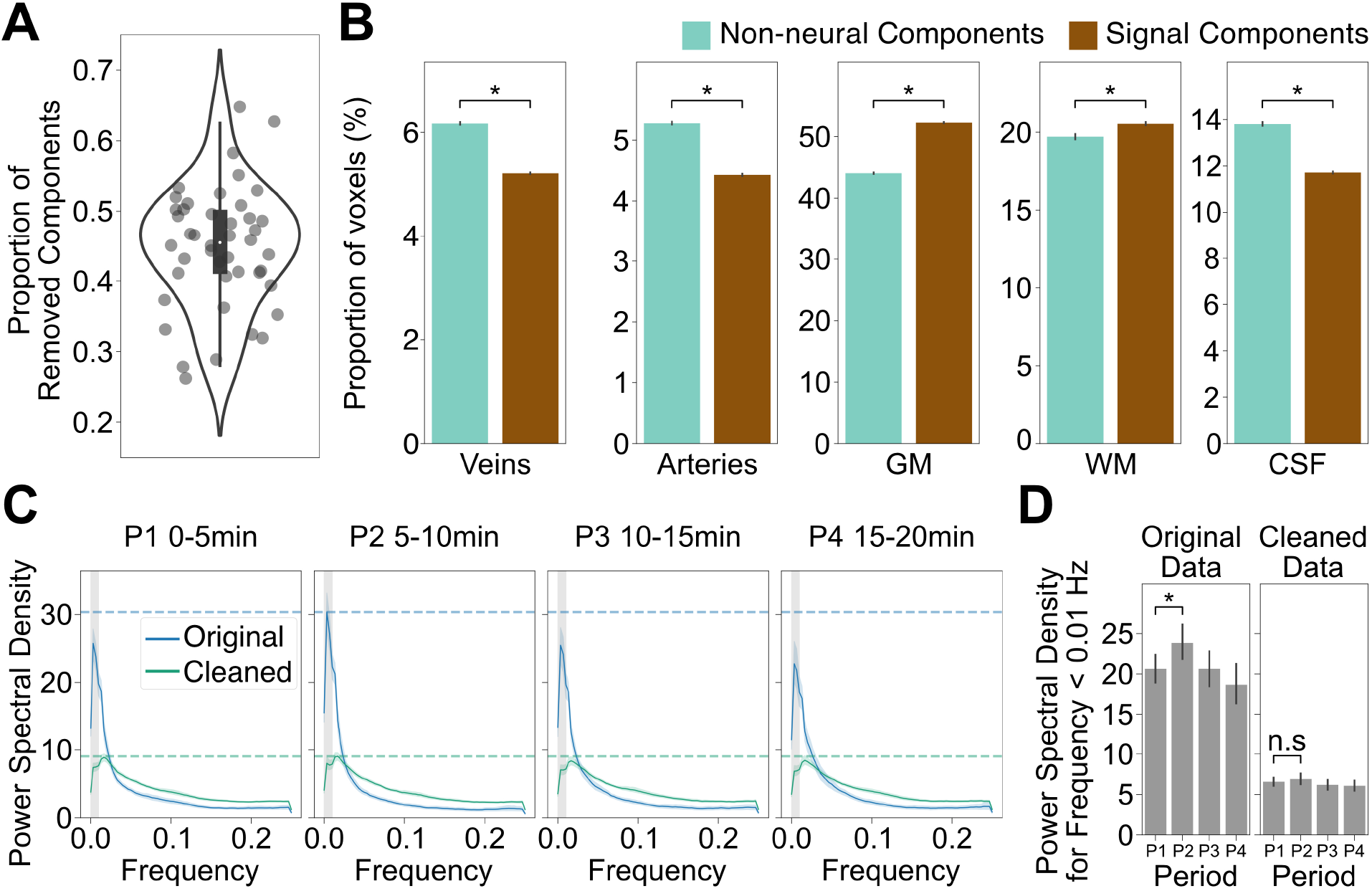
Characterization of artifactual components identified and removed by FIX-ICA. (A) Independent Component Analysis (ICA), combined with FMRIB’s ICA-based X-noiseifier (FIX), was applied to identify and remove non-neural components from each resting-state fMRI run. Each dot represents the proportion of components classified as artifacts for an individual run, and the violin plot illustrates the distribution across all runs. (B) The mean tissue probabilities (from vein, artery, GM, WM, and CSF tissue probability maps) were calculated for components classified as “signal” or “ non-neural.” Non-parametric bootstrappings estimate the differences in tissue probability between signal and non-neural components, revealing tissue-specific tendencies of artifactual components. *Testing significance by estimating 95% CI with multiple comparison. (C) Power spectral analysis was conducted for each 5-min phase, comparing data before and after the cleaning procedure. The original data exhibited elevated spectral power in the extremely low-frequency range (< 0.01 Hz; shaded area), which is commonly attributed to scanner drift, slow physiological fluctuations, or head motion. These non-neuronal fluctuations were largely removed by the FIX cleanup, resulting in reduced power in this frequency band in the cleaned data. (D) The increase in extremely low-frequency power during oxygen inhalation (P2) relative to baseline (P1) suggests that oxygen delivery induced additional low-frequency noise. After cleaning procedure, this difference was no longer present, indicating that these fluctuations were effectively identified as non-neuronal and removed, preserving more reliable BOLD signals for further analysis.

**Supplementary Figure 3.**
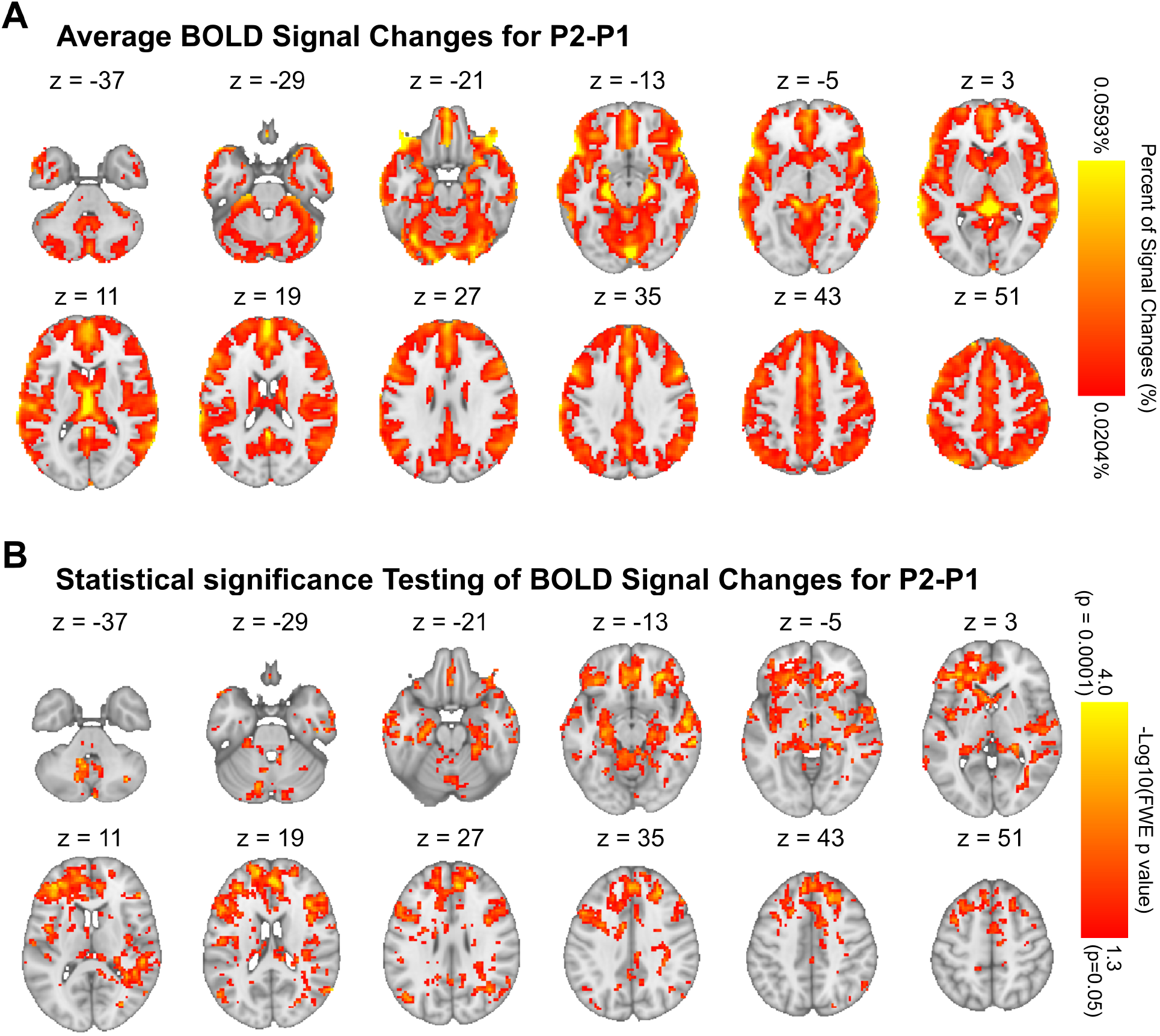
Voxel-wise map for BOLD signal changes (P2–P1) and statistical significance testing. (A) Voxel-wise map comparing oxygen inhalation (P2) to baseline (P1), displayed as percent signal changes. Using the cleaned data, signal changes were calculated and thresholded at 50% of the highest value (0.0204%) within the whole brain mask. The brightest color represents 98% of the highest signal value (0.0593%). With nearly all voxels exhibiting positive values, only positive signal changes are shown. (B) Statistical significance map. Non-parametric permutation testing was used to assess significance while controlling for multiple comparisons. The map is thresholded with voxel-level family-wise error (FWE) correction at p < 0.05, with the brightest color indicating voxel-level FWE p at 0.001.

**Supplementary Figure 4.**
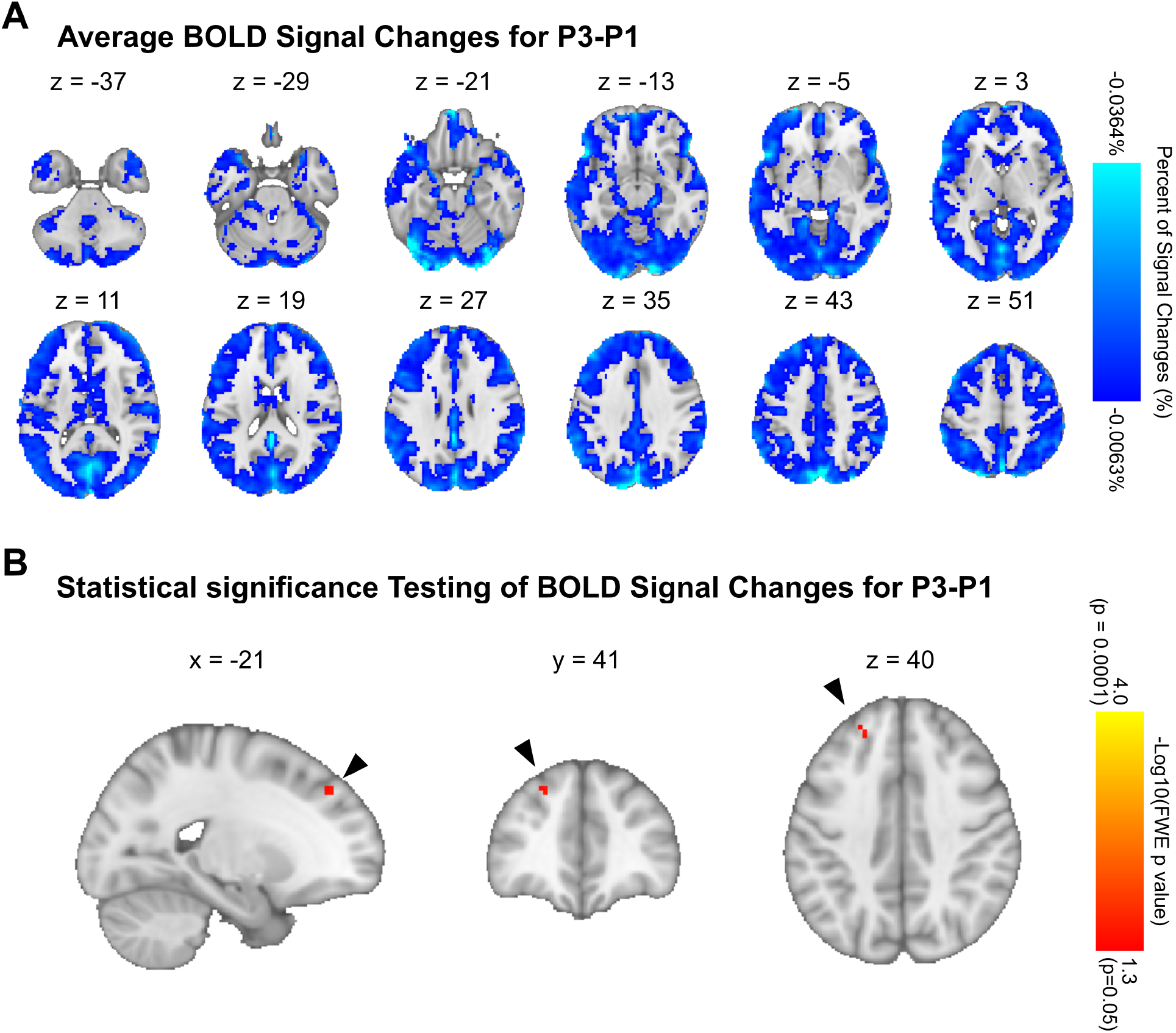
Voxel-wise map for BOLD signal changes (P3–P1) and statistical significance testing. (A) Voxelwise map comparing immediate oxygen withdrwal (P3) to baseline (P1), displayed as percent signal changes. Using the cleaned data, signal changes were calculated and thresholded at 50% of the lowest value (−0.0063%) within the whole brain mask. The brightest color represents 98% of the highest signal value (−0.0364%). With nearly all voxels exhibiting negative values, only negative signal changes are shown. (B) Statistical significance map. Non-parametric permutation testing was used to assess significance while controlling for multiple comparisons. The map is thresholded with voxel-level family-wise error (FWE) correction at p < 0.05, with the brightest color indicating voxel-level FWE p at 0.001.

**Supplementary Figure 5.**
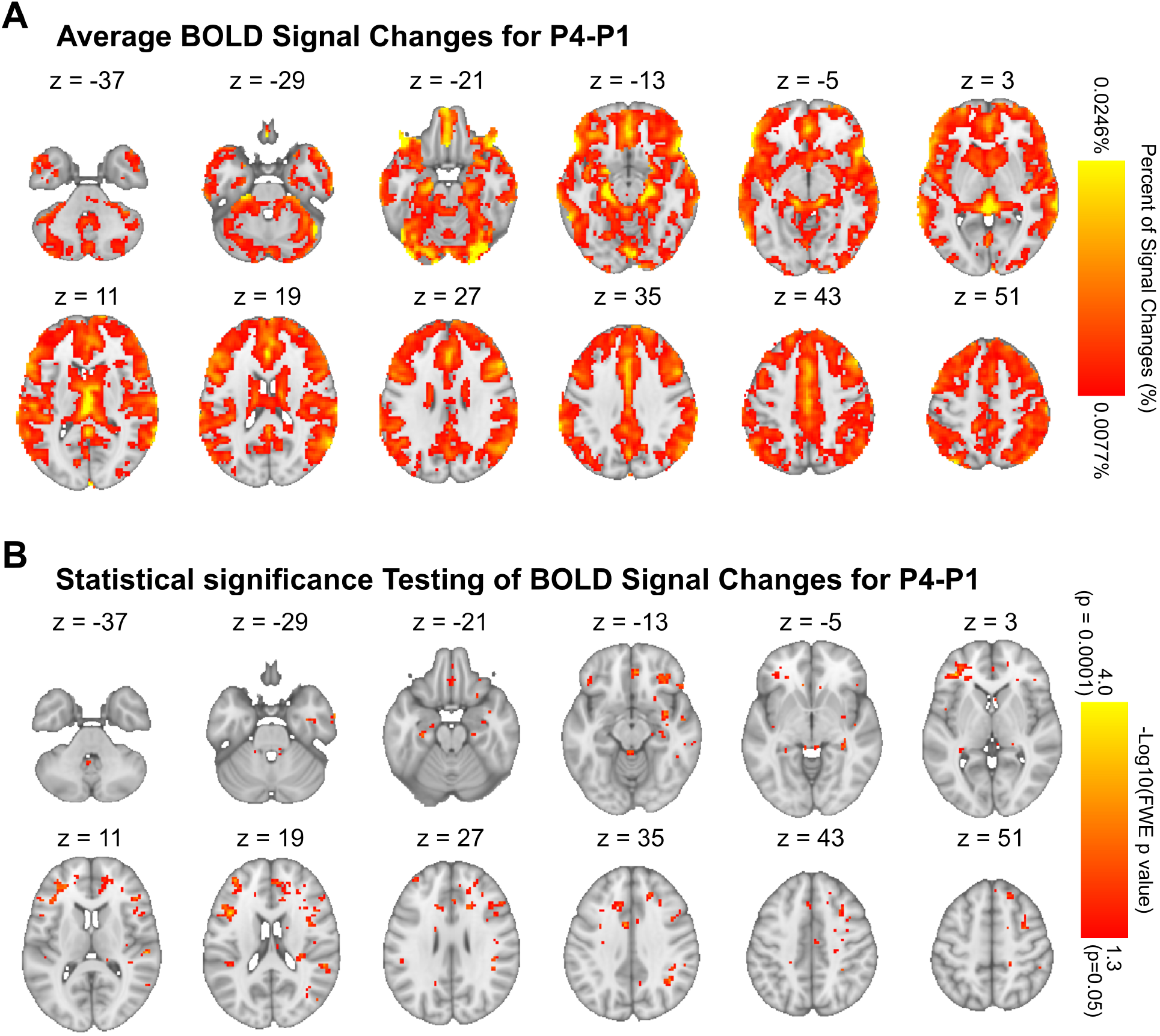
Voxel-wise map for BOLD signal changes (P4–P1) and statistical significance testing. (A) Voxelwise map comparing 5 min after oxygen withdrwal (P4) to baseline (P1), displayed as percent signal changes. Using the cleaned data, signal changes were calculated and thresholded at 50% of the highest value (0.0077%) within the whole brain mask. The brightest color represents 98% of the highest signal value (0.0246%). With nearly all voxels exhibiting positive values, only positive signal changes are shown. (B) Statistical significance map. Non-parametric permutation testing was used to assess significance while controlling for multiple comparisons. The map is thresholded with voxel-level family-wise error (FWE) correction at p < 0.05, with the brightest color indicating voxel-level FWE p at 0.001.

**Supplementary Figure 6.**
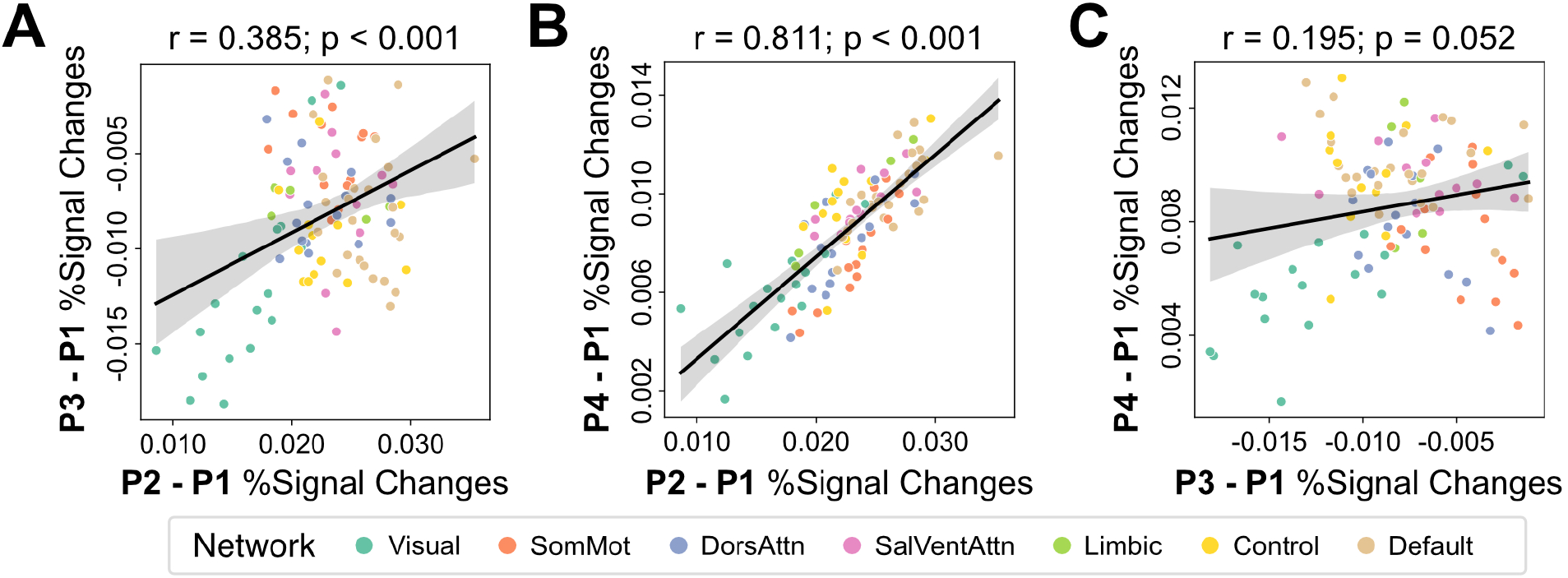
Correlations of signal changes across 100 cortical regions between different phases. Group-averaged signal changes from 100 cortical regions were extracted for different phases, and correlations of these changes were computed: (A)between P2–P1 and P3–P1, (B) between P2–P1 and P4–P1, and (C) between P3–P1 and P4–P1, to examine similarities and differences in response patterns across phases. Each dot represents a cortical region, with color indicating its associated network. All correlations are positive, with the highest correlation observed between P2–P1 and P4–P1. While these positive correlations suggest general trends in the response patterns, some deviations from these trends are evident across the different phases.

**Supplementary Figure 7.**
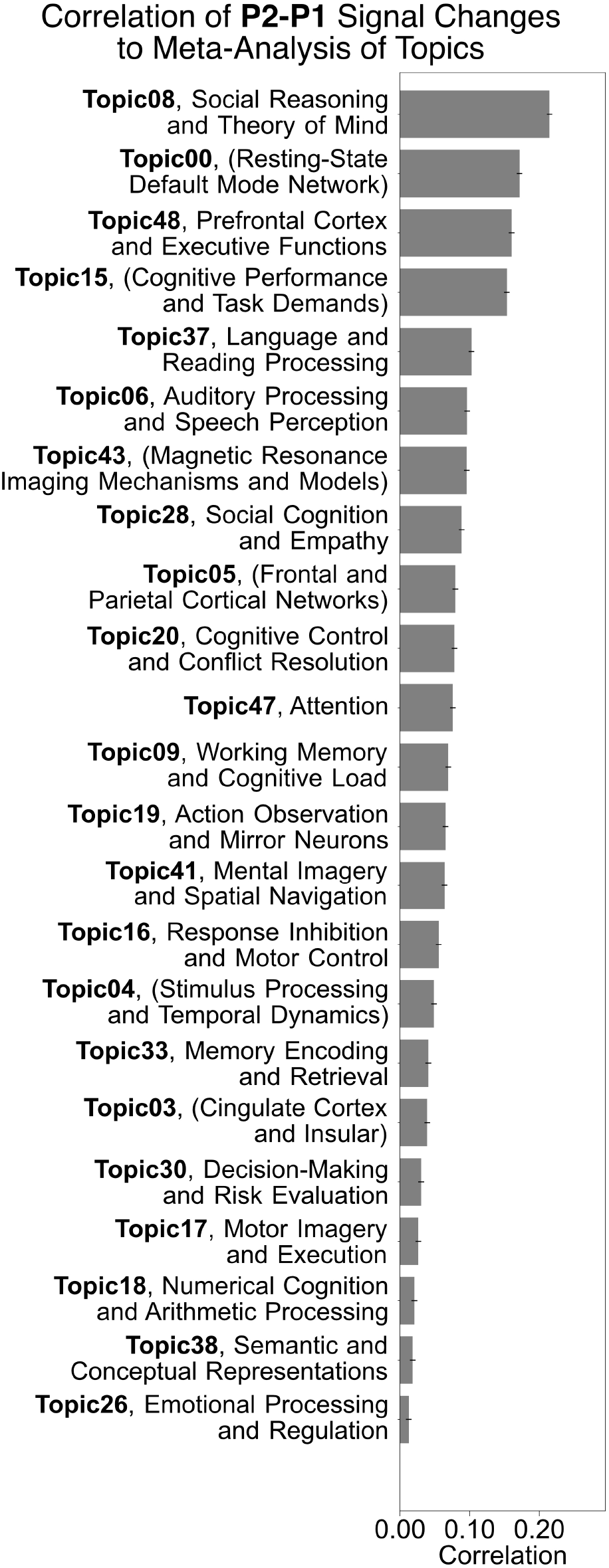
Functional decoding results for P2–P1 (oxygen inhalation). The voxel-wise signal change map of BOST_*p*2−*p*1_was submitted to functional decoding analysis to interpret how oxygen-induced BOLD signal changes relate to functional outcomes. The bars display the correlation coefficients, sorted from highest to lowest values. Topic labels were generated by ChatGPT from consistent terms and manually reviewed, and non-cognitive topics are marked in parentheses.

**Supplementary Figure 8.**
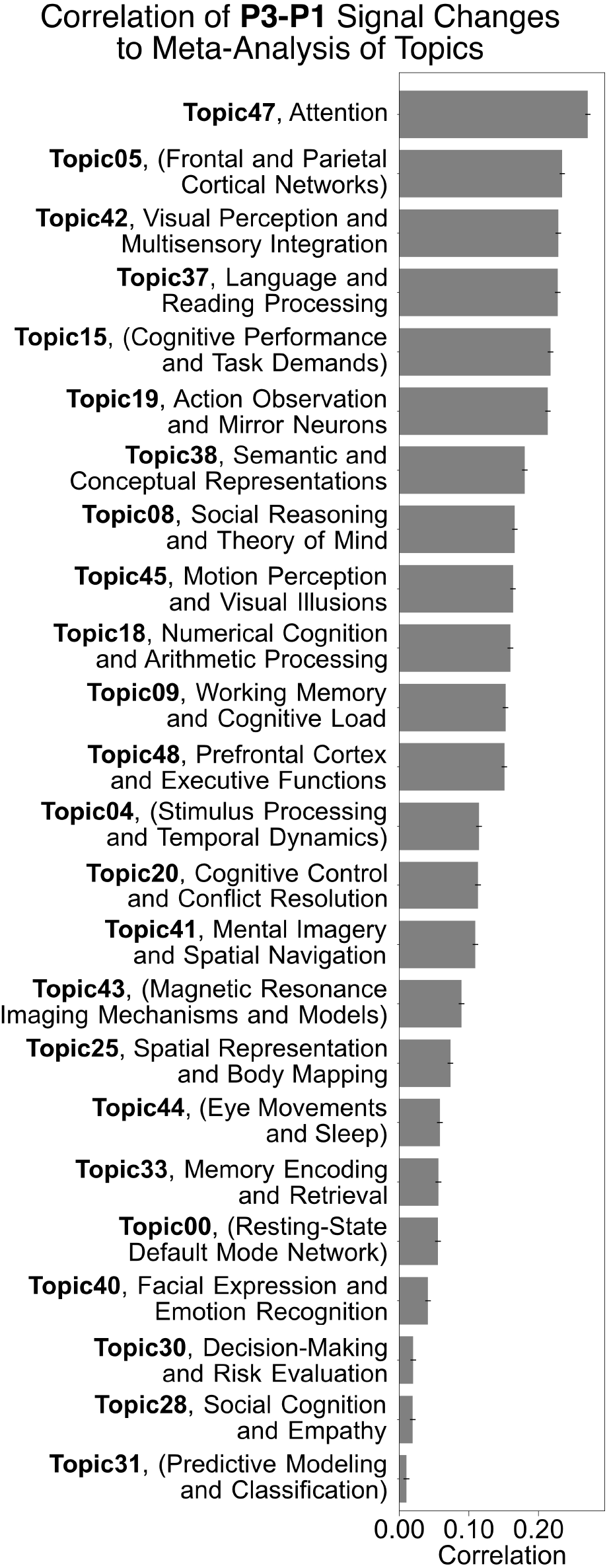
Functional decoding results for P3–P1 (immediate oxygen withdrawal). The voxel-wise signal change map of BOST_*p*3−*p*1_was submitted to functional decoding analysis to interpret how oxygen-induced BOLD signal changes relate to functional outcomes. The bars display the correlation coefficients, sorted from highest to lowest values. Topic labels were generated by ChatGPT from consistent terms and manually reviewed, and non-cognitive topics are marked in parentheses.

**Supplementary Figure 9.**
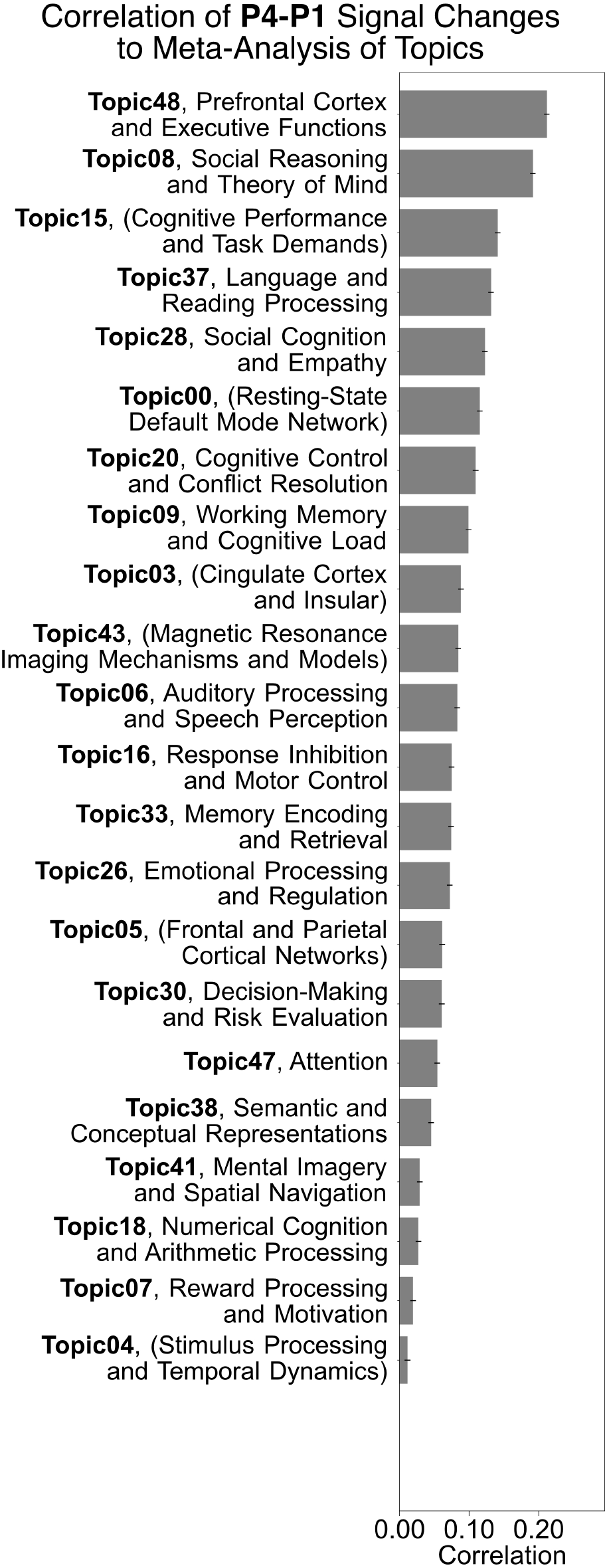
Functional decoding results for P4–P1 (post-oxygen rebound). The voxel-wise signal change map of BOST_*p*4−*p*1_was submitted to functional decoding analysis to interpret how oxygen-induced BOLD signal changes relate to functional outcomes. The bars display the correlation coefficients, sorted from highest to lowest values. Topic labels were generated by ChatGPT from consistent terms and manually reviewed, and non-cognitive topics are marked in parentheses.

## Bibliography

1. Pierre J. Magistretti and Igor Allaman. A Cellular Perspective on Brain Energy Metabolism and Functional Imaging. Neuron, 86(4):883–901, May 2015. ISSN 0896-6273. doi: 10.1016/j.neuron.2015.03.035.

2. Arnab Ghosh, David Highton, Christina Kolyva, Ilias Tachtsidis, Clare E Elwell, and Martin Smith. Hyperoxia results in increased aerobic metabolism following acute brain injury. Journal of Cerebral Blood Flow & Metabolism, 37(8):2910–2920, August 2017. ISSN 0271-678X. doi: 10.1177/0271678X16679171.

3. Gauri H. Malthankar-Phatak, Anant B. Patel, Ying Xia, Soonsun Hong, Golam M. I. Chowdhury, Kevin L. Behar, Isaac A. Orina, and James C. K. Lai. Effects of continuous hypoxia on energy metabolism in cultured cerebro-cortical neurons. Brain Research, 1229:147–154, September 2008. ISSN 0006-8993. doi: 10.1016/j.brainres.2008.06.074.

4. David M Shaw, Peter M Bloomfield, and Nicholas Gant. The effect of acute normobaric hyperoxia on cognition: A systematic review, meta-analysis and meta-regression. Physiology & Behavior, 267:114208, August 2023. ISSN 00319384. doi: 10.1016/j.physbeh.2023.114208.

5. Anna B. Marcinkowska, Natalia D. Mankowska, Jacek Kot, and Pawel J. Winklewski. Impact of Hyperbaric Oxygen Therapy on Cognitive Functions: A Systematic Review. Neuropsychology Review, 32(1):99–126, March 2022. ISSN 1573-6660. doi: 10.1007/s11065-021-09500-9.

6. Amir Hadanny, Malka Daniel-Kotovsky, Gil Suzin, Rahav Boussi-Gross, Merav Catalogna, Kobi Dagan, Yafit Hachmo, Ramzia Abu Hamed, Efrat Sasson, Gregory Fishlev, Erez Lang, Nir Polak, Keren Doenyas, Mony Friedman, Sigal Tal, Yonatan Zemel, Yair Bechor, and Shai Efrati. Cognitive enhancement of healthy older adults using hyperbaric oxygen: A randomized controlled trial. Aging, 12(13):13740–13761, June 2020. ISSN 1945-4589. doi: 10.18632/aging.103571.

7. Priya Balasubramanian, Jordan Delfavero, Adam Nyul-Toth, Amber Tarantini, Rafal Gulej, and Stefano Tarantini. Integrative Role of Hyperbaric Oxygen Therapy on Healthspan, Age-Related Vascular Cognitive Impairment, and Dementia. Frontiers in Aging, 2, September 2021. ISSN 2673-6217. doi: 10.3389/fragi.2021.678543.

8. Soon-Cheol Chung, Ji-Hun Kwon, Hang-Woon Lee, Gye-Rae Tack, Bongsoo Lee, Jeong-Han Yi, and Soo-Yeol Lee. Effects of high concentration oxygen administration on n −back task performance and physiological signals. Physiological Measurement, 28(4):389–396, April 2007. ISSN 0967-3334, 1361-6579. doi: 10.1088/0967-3334/28/4/005.

9. Andrew B Scholey, Mark C Moss, Nick Neave, and Keith Wesnes. Cognitive Performance, Hyperoxia, and Heart Rate Following Oxygen Administration in Healthy Young Adults. Physiology & Behavior, 67(5):783–789, November 1999. ISSN 0031-9384. doi: 10.1016/S0031-9384(99)00183-3.

10. Sayeed A. D. Kizuk, Wesley Vuong, Joanna E. MacLean, Clayton T. Dickson, and Kyle E. Mathewson. Electrophysiological correlates of hyperoxia during resting-state EEG in awake human subjects. Psychophysiology, 56(10):e13401, 2019. ISSN 1469-8986. doi: 10.1111/psyp.13401.

11. Ho-Jun Seo, Won-Myong Bahk, Tae-Yun Jun, and Jeong-Ho Chae. The Effect of Oxygen Inhalation on Cognitive Function and EEG in Healthy Adults. Clinical Psychopharmacology and Neuroscience, 5(1):25–30, February 2007. ISSN 1738-1088.

12. Min Sheng, Peiying Liu, Deng Mao, Yulin Ge, and Hanzhang Lu. The impact of hyperoxia on brain activity: A resting-state and task-evoked electroencephalography (EEG) study. PLOS ONE, 12(5): e0176610, 2017. ISSN 1932-6203. doi: 10.1371/journal.pone.0176610.

13. KK Kwong, JW Belliveau, D. Chesler, IE Goldberg, RM Weisskoff, BP Poncelet, DN Kennedy, BE Hoppel, MS Cohen, and R Turner. Dynamic magnetic resonance imaging of human brain activity during primary sensory stimulation. Proceedings of the National Academy of Sciences, 89(12):5675–5679, June 1992. doi: 10.1073/pnas.89.12.5675.

14. S Ogawa, DW Tank, R Menon, JM Ellermann, SG Kim, H Merkle, and K Ugurbil. Intrinsic signal changes accompanying sensory stimulation: Functional brain mapping with magnetic resonance imaging. Proceedings of the National Academy of Sciences, 89(13):5951–5955, July 1992. doi: 10.1073/pnas.89.13.5951.

15. Richard B Buxton. The physics of functional magnetic resonance imaging (fMRI). Reports on Progress in Physics, 76(9):096601, September 2013. ISSN 0034-4885. doi: 10.1088/0034-4885/76/9/096601.

16. Richard B. Buxton, Kâmil Uludağ, David J. Dubowitz, and Thomas T. Liu. Modeling the hemodynamic response to brain activation. NeuroImage, 23:S220–S233, January 2004. ISSN 1053-8119. doi: 10.1016/j.neuroimage.2004.07.013.

17. Nicholas P. Blockley, Valerie E. M. Griffeth, Michael A. Germuska, Daniel P. Bulte, and Richard B. Buxton. An analysis of the use of hyperoxia for measuring venous cerebral blood volume: Comparison of the existing method with a new analysis approach. NeuroImage, 72:33–40, May 2013. ISSN 1053-8119. doi: 10.1016/j.neuroimage.2013.01.039.

18. Peter A. Chiarelli, Daniel P. Bulte, Richard Wise, Daniel Gallichan, and Peter Jezzard. A calibration method for quantitative BOLD fMRI based on hyperoxia. NeuroImage, 37(3):808–820, September 2007. ISSN 1053-8119. doi: 10.1016/j.neuroimage.2007.05.033.

19. Yuhan Ma, Avery J.L. Berman, and G. Bruce Pike. The effect of dissolved oxygen on the relaxation rates of blood plasma: Implications for hyperoxia calibrated BOLD. Magnetic Resonance in Medicine, 76(6):1905–1911, 2016. ISSN 1522-2594. doi: 10.1002/mrm.26069.

20. Christoph Losert, Michael Peller, Philipp Schneider, and Maximilian Reiser. Oxygen-enhanced MRI of the brain. Magnetic Resonance in Medicine, 48(2):271–277, 2002. ISSN 1522-2594. doi: 10.1002/mrm.10215.

21. Georgia Hardavella, Ioannis Karampinis, Armin Frille, Katherina Sreter, and Ilona Rousalova. Oxygen devices and delivery systems. Breathe, 15(3):e108–e116, October 2019. ISSN 1810-6838, 2073-4735. doi: 10.1183/20734735.0204-2019.

22. Damon P. Cardenas, Eric R. Muir, Shiliang Huang, Angela Boley, Daniel Lodge, and Timothy Q. Duong. Functional MRI during hyperbaric oxygen: Effects of oxygen on neurovascular coupling and BOLD fMRI signals. NeuroImage, 119:382–389, October 2015. ISSN 1053-8119. doi: 10.1016/j.neuroimage.2015.06.082.

23. Elizabeth G. Damato, Tod A. Flak, Ryan S. Mayes, Kingman P. Strohl, Aemilee M. Ziganti, Alireza Abdollahifar, Chris A. Flask, Joseph C. LaManna, and Michael J. Decker. Neurovascular and cortical responses to hyperoxia: Enhanced cognition and electroencephalographic activity despite reduced perfusion. The Journal of Physiology, 598(18):3941–3956, 2020. ISSN 1469-7793. doi: 10.1113/JP279453.

24. Andrew B. Scholey, Sarah Benson, Shirley Sela-Venter, Marlou Mackus, and Mark C. Moss. Oxygen Administration and Acute Human Cognitive Enhancement: Higher Cognitive Demand Leads to a More Rapid Decay of Transient Hyperoxia. Journal of Cognitive Enhancement, 4(1):94–99, March 2020. ISSN 2509-3290, 2509-3304. doi: 10.1007/s41465-019-00145-4.

25. Yoann Gole, Ombeline Gargne, Mathieu Coulange, Jean-Guillaume Steinberg, Malika Bouhaddi, Yves Jammes, Jacques Regnard, and Alain Boussuges. Hyperoxia-induced alterations in cardiovascular function and autonomic control during return to normoxic breathing. European Journal of Applied Physiology, 111(6):937–946, June 2011. ISSN 1439-6327. doi: 10.1007/s00421-010-1711-4.

26. Michaël Bernier, Stephen C. Cunnane, and Kevin Whittingstall. The morphology of the human cerebrovascular system. Human Brain Mapping, 39(12):4962–4975, 2018. ISSN 1097-0193. doi: 10.1002/hbm.24337.

27. Alexander Schaefer, Ru Kong, Evan M Gordon, Timothy O Laumann, Xi-Nian Zuo, Avram J Holmes, Simon B Eickhoff, and BT Thomas Yeo. Local-Global Parcellation of the Human Cerebral Cortex from Intrinsic Functional Connectivity MRI. Cerebral Cortex, 28(9):3095–3114, September 2018. ISSN 1047-3211. doi: 10.1093/cercor/bhx179.

28. Ludovica Griffanti, Gwenaëlle Douaud, Janine Bijsterbosch, Stefania Evangelisti, Fidel Alfaro-Almagro, Matthew F. Glasser, Eugene P. Duff, Sean Fitzgibbon, Robert Westphal, Davide Carone, Christian F. Beckmann, and Stephen M. Smith. Hand classification of fMRI ICA noise components. NeuroImage, 154:188–205, July 2017. ISSN 1053-8119. doi: 10.1016/j.neuroimage.2016.12.036.

29. Ludovica Griffanti, Gholamreza Salimi-Khorshidi, Christian F. Beckmann, Edward J. Auerbach, Gwenaëlle Douaud, Claire E. Sexton, Enikő Zsoldos Klaus P. Ebmeier, Nicola Filippini, Clare E. Mackay, Steen Moeller, Junqian Xu, Essa Yacoub, Giuseppe Baselli, Kamil Ugurbil, Karla L. Miller, and Stephen M. Smith. ICA-based artefact removal and accelerated fMRI acquisition for improved resting state network imaging. NeuroImage, 95:232–247, July 2014. ISSN 1053-8119. doi: 10.1016/j.neuroimage.2014.03.034.

30. Gholamreza Salimi-Khorshidi, Gwenaëlle Douaud, Christian F. Beckmann, Matthew F. Glasser, Ludovica Griffanti, and Stephen M. Smith. Automatic denoising of functional MRI data: Combining independent component analysis and hierarchical fusion of classifiers. NeuroImage, 90:449–468, April 2014. ISSN 1053-8119. doi: 10.1016/j.neuroimage.2013.11.046.

31. Marta Bianciardi, Masaki Fukunaga, Peter van Gelderen, Silvina G. Horovitz, Jacco A. de Zwart, Karin Shmueli, and Jeff H. Duyn. Sources of functional magnetic resonance imaging signal fluctuations in the human brain at rest: A 7 T study. Magnetic Resonance Imaging, 27(8):1019–1029, October 2009. ISSN 0730-725X. doi: 10.1016/j.mri.2009.02.004.

32. Tal Yarkoni, Russell A. Poldrack, Thomas E. Nichols, David C. Van Essen, and Tor D. Wager. Large-scale automated synthesis of human functional neuroimaging data. Nature Methods, 8(8): 665–670, August 2011. ISSN 1548-7105. doi: 10.1038/nmeth.1635.

33. Andrew Zalesky, Alex Fornito, and Edward T. Bullmore. Network-based statistic: Identifying differences in brain networks. NeuroImage, 53(4):1197–1207, December 2010. ISSN 1053-8119. doi: 10.1016/j.neuroimage.2010.06.041.

34. David Attwell and Simon B. Laughlin. An Energy Budget for Signaling in the Grey Matter of the Brain. Journal of Cerebral Blood Flow & Metabolism, 21(10):1133–1145, October 2001. ISSN 0271-678X. doi: 10.1097/00004647-200110000-00001.

35. Giuseppina Giannì, Andrea Minini, Sara Fratino, Lorenzo Peluso, Filippo Annoni, Mauro Oddo, Sophie Schuind, Jacques Creteur, Fabio Silvio Taccone, and Elisa Gouvêa Bogossian. The Impact of Short-Term Hyperoxia on Cerebral Metabolism: A Systematic Review and Meta-Analysis. Neurocritical Care, 37(2):547–557, October 2022. ISSN 1556-0961. doi: 10.1007/s12028-022-01529-9.

36. Daniel P Bulte, Peter A Chiarelli, Richard G Wise, and Peter Jezzard. Cerebral Perfusion Response to Hyperoxia. Journal of Cerebral Blood Flow & Metabolism, 27(1):69–75, January 2007. ISSN 0271-678X. doi: 10.1038/sj.jcbfm.9600319.

37. Thomas F. Floyd, James M. Clark, Robert Gelfand, John A. Detre, Sarah Ratcliffe, Dimitri Guvakov, Christian J. Lambertsen, and Roderic G. Eckenhoff. Independent cerebral vasoconstrictive effects of hyperoxia and accompanying arterial hypocapnia at 1 ATA. Journal of Applied Physiology, 95(6):2453–2461, December 2003. ISSN 8750-7587. doi: 10.1152/japplphysiol.00303.2003.

38. Yuhan Ma, Erin L. Mazerolle, Junghun Cho, Hongfu Sun, Yi Wang, and G. Bruce Pike. Quantification of brain oxygen extraction fraction using QSM and a hyperoxic challenge. Magnetic Resonance in Medicine, 84(6):3271–3285, 2020. ISSN 1522-2594. doi: 10.1002/mrm.28390.

39. Gabriella MK Rossetti, Giovanni d’Avossa, Matthew Rogan, Jamie H Macdonald, Samuel J Oliver, and Paul G Mullins. Reversal of neurovascular coupling in the default mode network: Evidence from hypoxia. Journal of Cerebral Blood Flow & Metabolism, 41(4):805–818, April 2021. ISSN 0271-678X. doi: 10.1177/0271678X20930827.

40. Kenji Ishibashi, Keita Sakurai, Keigo Shimoji, Aya M. Tokumaru, and Kenji Ishii. Altered functional connectivity of the default mode network by glucose loading in young, healthy participants. BMC Neuroscience, 19(1):33, May 2018. ISSN 1471-2202. doi: 10.1186/s12868-018-0433-0.

41. Tyler Blazey, Abraham Z Snyder, Yi Su, Manu S Goyal, John J Lee, Andrei G Vlassenko, Ana Maria Arbeláez, and Marcus E Raichle. Quantitative positron emission tomography reveals regional differences in aerobic glycolysis within the human brain. Journal of Cerebral Blood Flow & Metabolism, 39(10):2096–2102, October 2019. ISSN 0271-678X. doi: 10.1177/0271678X18767005.

42. S. Neil Vaishnavi, Andrei G. Vlassenko, Melissa M. Rundle, Abraham Z. Snyder, Mark A. Mintun, and Marcus E. Raichle. Regional aerobic glycolysis in the human brain. Proceedings of the National Academy of Sciences, 107(41):17757–17762, October 2010. doi: 10.1073/pnas.1010459107.

43. Fahmeed Hyder, Robert K Fulbright, Robert G Shulman, and Douglas L Rothman. Glutamatergic Function in the Resting Awake Human Brain is Supported by Uniformly High Oxidative Energy. Journal of Cerebral Blood Flow & Metabolism, 33(3):339–347, March 2013. ISSN 0271-678X. doi: 10.1038/jcbfm.2012.207.

44. Fahmeed Hyder, Peter Herman, Christopher J Bailey, Arne Møller, Ronen Globinsky, Robert K Fulbright, Douglas L Rothman, and Albert Gjedde. Uniform distributions of glucose oxidation and oxygen extraction in gray matter of normal human brain: No evidence of regional differences of aerobic glycolysis. Journal of Cerebral Blood Flow & Metabolism, 36(5):903–916, May 2016. ISSN 0271-678X. doi: 10.1177/0271678X15625349.

45. Gregg L. Semenza. Hypoxia-Inducible Factors in Physiology and Medicine. Cell, 148(3):399–408, February 2012. ISSN 0092-8674. doi: 10.1016/j.cell.2012.01.021.

46. Angeles Almeida, Julia Almeida, Juan P. Bolaños, and Salvador Moncada. Different responses of astrocytes and neurons to nitric oxide: The role of glycolytically generated ATP in astrocyte protection. Proceedings of the National Academy of Sciences, 98(26):15294–15299, December 2001. doi: 10.1073/pnas.261560998.

47. Guy C. Brown and Chris.E. Cooper. Nanomolar concentrations of nitric oxide reversibly inhibit synaptosomal respiration by competing with oxygen at cytochrome oxidase. FEBS Letters, 356(2-3): 295–298, 1994. ISSN 1873-3468. doi: 10.1016/0014-5793(94)01290-3.

48. Maria Erecińska, David Nelson, and Jane M. Vanderkooi. Effects of NO-Generating Compounds on Synaptosomal Energy Metabolism. Journal of Neurochemistry, 65(6):2699–2705, 1995. ISSN 1471-4159. doi: 10.1046/j.1471-4159.1995.65062699.x.

49. G. M. Rubanyi and P. M. Vanhoutte. Superoxide anions and hyperoxia inactivate endothelium-derived relaxing factor. American Journal of Physiology-Heart and Circulatory Physiology, 250(5):H822–H827, May 1986. ISSN 0363-6135. doi: 10.1152/ajpheart.1986.250.5.H822.

50. Darko Modun, Mladen Krnic, Jonatan Vukovic, Visnja Kokic, Lea Kukoc-Modun, Dimitrios Tsikas, and Zeljko Dujic. Plasma nitrite concentration decreases after hyperoxia-induced oxidative stress in healthy humans. Clinical Physiology and Functional Imaging, 32(5):404–408, 2012. ISSN 1475-097X. doi: 10.1111/j.1475-097X.2012.01133.x.

51. Ranjithmenon Muraleedharan, Mruniya V. Gawali, Durgesh Tiwari, Abitha Sukumaran, Nicole Oatman, Jane Anderson, Diana Nardini, Mohammad Alfrad Nobel Bhuiyan, Ivan Tkáč, Amber Lynne Ward, Mondira Kundu, Ronald Waclaw, Lionel M. Chow, Christina Gross, Raghavendra Rao, Stefanie Schirmeier, and Biplab Dasgupta. AMPK-Regulated Astrocytic Lactate Shuttle Plays a Non-Cell-Autonomous Role in Neuronal Survival. Cell Reports, 32(9), September 2020. ISSN 2211-1247. doi: 10.1016/j.celrep.2020.108092.

52. Christian Pehmøller, Jonas T. Treebak, Jesper B. Birk, Shuai Chen, Carol MacKintosh, D. Grahame Hardie, Erik A. Richter, and Jørgen F. P. Wojtaszewski. Genetic disruption of AMPK signaling abolishes both contraction- and insulin-stimulated TBC1D1 phosphorylation and 14-3-3 binding in mouse skeletal muscle. American Journal of Physiology-Endocrinology and Metabolism, 297(3): E665–E675, September 2009. ISSN 0193-1849. doi: 10.1152/ajpendo.00115.2009.

53. Ning Wu, Bin Zheng, Adam Shaywitz, Yossi Dagon, Christine Tower, Gary Bellinger, Che-Hung Shen, Jennifer Wen, John Asara, Timothy E. McGraw, Barbara B. Kahn, and Lewis C. Cantley. AMPK-Dependent Degradation of TXNIP upon Energy Stress Leads to Enhanced Glucose Uptake via GLUT1. Molecular Cell, 49(6):1167–1175, March 2013. ISSN 1097-2765. doi: 10.1016/j.molcel.2013.01.035.

54. Sean L. McGee, Bryce J.W. van Denderen, Kirsten F. Howlett, Janelle Mollica, Jonathan D. Schertzer, Bruce E. Kemp, and Mark Hargreaves. AMP-Activated Protein Kinase Regulates GLUT4 Transcription by Phosphorylating Histone Deacetylase 5. Diabetes, 57(4):860–867, April 2008. ISSN 0012-1797. doi: 10.2337/db07-0843.

55. Richard M. Reznick and Gerald I. Shulman. The role of AMP-activated protein kinase in mitochondrial biogenesis. The Journal of Physiology, 574(1):33–39, 2006. ISSN 1469-7793. doi: 10.1113/jphysiol.2006.109512.

56. Sławomir Kujawski, Joanna Słomko, Karl J. Morten, Modra Murovska, Katarzyna Buszko, Julia L. Newton, and Paweł Zalewski. Autonomic and Cognitive Function Response to Normobaric Hyperoxia Exposure in Healthy Subjects. Preliminary Study. Medicina, 56(4):172, April 2020. ISSN 1648-9144. doi: 10.3390/medicina56040172.

57. Ronghao Yu, Bin Wang, Shumei Li, Junjing Wang, Feng Zhou, Shufang Chu, Xianyou He, Xue Wen, Xiaoxiao Ni, Liqing Liu, Qiuyou Xie, and Ruiwang Huang. Cognitive enhancement of healthy young adults with hyperbaric oxygen: A preliminary resting-state fMRI study. Clinical Neurophysiology, 126(11):2058–2067, November 2015. ISSN 1388-2457. doi: 10.1016/j.clinph.2015.01.010.

58. Rong Wang, Mianxin Liu, Xinhong Cheng, Ying Wu, Andrea Hildebrandt, and Changsong Zhou. Segregation, integration, and balance of large-scale resting brain networks configure different cognitive abilities. Proceedings of the National Academy of Sciences, 118(23):e2022288118, June 2021. doi: 10.1073/pnas.2022288118.

59. Carissa L. Philippi, Daniel Tranel, Melissa Duff, and David Rudrauf. Damage to the default mode network disrupts autobiographical memory retrieval. Social Cognitive and Affective Neuroscience, 10(3):318–326, March 2015. ISSN 1749-5016. doi: 10.1093/scan/nsu070.

60. R. Nathan Spreng, Raymond A. Mar, and Alice S. N. Kim. The Common Neural Basis of Autobiographical Memory, Prospection, Navigation, Theory of Mind, and the Default Mode: A Quantitative Meta-analysis. Journal of Cognitive Neuroscience, 21(3):489–510, March 2009. ISSN 0898-929X. doi: 10.1162/jocn.2008.21029.

61. Ben Shofty, Tal Gonen, Eyal Bergmann, Naama Mayseless, Akiva Korn, Simone Shamay-Tsoory, Rachel Grossman, Itamar Jalon, Itamar Kahn, and Zvi Ram. The default network is causally linked to creative thinking. Molecular Psychiatry, 27(3):1848–1854, March 2022. ISSN 1476-5578. doi: 10.1038/s41380-021-01403-8.

62. Paul M Macey, Katherine E Macey, Rajesh Kumar, and Ronald M Harper. A method for removal of global effects from fMRI time series. NeuroImage, 22(1):360–366, May 2004. ISSN 1053-8119. doi: 10.1016/j.neuroimage.2003.12.042.

63. Marta Bianciardi, Masaki Fukunaga, Peter van Gelderen, Jacco A de Zwart, and Jeff H Duyn. Negative BOLD-fMRI Signals in Large Cerebral Veins. Journal of Cerebral Blood Flow & Metabolism, 31(2):401–412, February 2011. ISSN 0271-678X. doi: 10.1038/jcbfm.2010.164.

64. Andrew T. Curtis, R. Matthew Hutchison, and Ravi S. Menon. Phase based venous suppression in resting-state BOLD GE-fMRI. NeuroImage, 100:51–59, October 2014. ISSN 1053-8119. doi: 10.1016/j.neuroimage.2014.05.079.

65. Julia Huck, Anna-Thekla Jäger, Uta Schneider, Sophia Grahl, Audrey P. Fan, Christine Tardif, Arno Villringer, Pierre-Louis Bazin, Christopher J. Steele, and Claudine J. Gauthier. Modeling venous bias in resting state functional MRI metrics. Human Brain Mapping, 44(14):4938–4955, 2023. ISSN 1097-0193. doi: 10.1002/hbm.26431.

66. Petra Ritter and Arno Villringer. Simultaneous EEG–fMRI. Neuroscience & Biobehavioral Reviews, 30(6):823–838, January 2006. ISSN 0149-7634. doi: 10.1016/j.neubiorev.2006.06.008.

67. Timothy L. Davis, Kenneth K. Kwong, Robert M. Weisskoff, and Bruce R. Rosen. Calibrated functional MRI: Mapping the dynamics of oxidative metabolism. Proceedings of the National Academy of Sciences, 95(4):1834–1839, February 1998. doi: 10.1073/pnas.95.4.1834.

68. Sungho Tak, Jonathan R. Polimeni, Danny J.J. Wang, Lirong Yan, and J. Jean Chen. Associations of Resting-State fMRI Functional Connectivity with Flow-BOLD Coupling and Regional Vasculature. Brain Connectivity, 5(3):137–146, April 2015. ISSN 2158-0014. doi: 10.1089/brain.2014.0299.

69. Lars Jonasson Stiernman, Filip Grill, Andreas Hahn, Lucas Rischka, Rupert Lanzenberger, Vania Panes Lundmark, Katrine Riklund, Jan Axelsson, and Anna Rieckmann. Dissociations between glucose metabolism and blood oxygenation in the human default mode network revealed by simultaneous PET-fMRI. Proceedings of the National Academy of Sciences, 118(27):e2021913118, July 2021. doi: 10.1073/pnas.2021913118.

70. Viviana Siless, Nicholas A. Hubbard, Robert Jones, Jonathan Wang, Nicole Lo, Clemens C. C. Bauer, Mathias Goncalves, Isabelle Frosch, Daniel Norton, Genesis Vergara, Kristina Conroy, Flavia Vaz De Souza, Isabelle M. Rosso, Aleena Hay Wickham, Elizabeth Ann Cosby, Megan Pinaire, Dina Hirshfeld-Becker, Diego A. Pizzagalli, Aude Henin, Stefan G. Hofmann, Randy P. Auerbach, Satrajit Ghosh, John Gabrieli, Susan Whitfield-Gabrieli, and Anastasia Yendiki. Image acquisition and quality assurance in the Boston Adolescent Neuroimaging of Depression and Anxiety study. NeuroImage: Clinical, 26:102242, January 2020. ISSN 2213-1582. doi: 10.1016/j.nicl.2020.102242.

71. Vladimir Fonov, Alan C. Evans, Kelly Botteron, C. Robert Almli, Robert C. McKinstry, and D. Louis Collins. Unbiased average age-appropriate atlases for pediatric studies. NeuroImage, 54(1): 313–327, January 2011. ISSN 1053-8119. doi: 10.1016/j.neuroimage.2010.07.033.

72. Bharat Biswal, F. Zerrin Yetkin, Victor M. Haughton, and James S. Hyde. Functional connectivity in the motor cortex of resting human brain using echo-planar mri. Magnetic Resonance in Medicine, 34(4):537–541, 1995. ISSN 1522-2594. doi: 10.1002/mrm.1910340409.

73. Rasmus M. Birn, Jason B. Diamond, Monica A. Smith, and Peter A. Bandettini. Separating respiratory-variation-related fluctuations from neuronal-activity-related fluctuations in fMRI. NeuroImage, 31(4):1536–1548, July 2006. ISSN 1053-8119. doi: 10.1016/j.neuroimage.2006.02.048.

74. Xi-Nian Zuo, Adriana Di Martino, Clare Kelly, Zarrar E. Shehzad, Dylan G. Gee, Donald F. Klein, F. Xavier Castellanos, Bharat B. Biswal, and Michael P. Milham. The oscillating brain: Complex and reliable. NeuroImage, 49(2):1432–1445, January 2010. ISSN 1053-8119. doi: 10.1016/j.neuroimage.2009.09.037.

75. Richard G Wise, Kojiro Ide, Marc J Poulin, and Irene Tracey. Resting fluctuations in arterial carbon dioxide induce significant low frequency variations in BOLD signal. NeuroImage, 21(4):1652–1664, April 2004. ISSN 1053-8119. doi: 10.1016/j.neuroimage.2003.11.025.

